# Murine Toll-like receptor 8 is a functional sensor of nucleic acid fragments

**DOI:** 10.1101/2025.10.14.682277

**Authors:** Lukas Hinkelmann, Erandi Rupasinghe, Mariam Brehm, Daniel S. Simpson, W. Samantha N. Jayasekara, Sunil Sapkota, Qihao Sun, Gunnar Kleinau, Victor Kumbol, Hugo McGurran, Ruitao Jin, Sitong He, Josiah Bones, Ana Kitanovic, Cassandra R. Harapas, Christina Krüger, Alexandra L. McAllan, Paul Borkenhagen, ChakYin Chan, Douglas Golenbock, Patrick Scheerer, Gerhard Krönke, Ben Corry, Lena Alexopoulou, Julia I. Ellyard, Eicke Latz, Stefan Bauer, Michael P. Gantier, Seija Lehnardt

## Abstract

Toll-like receptors 7 and 8 (TLR7 and TLR8) detect nucleic acid degradation products to initiate innate immune responses, although their ligand-sensing functions differ between species. Whereas human TLR8 is a well-established nucleic acid sensor, murine TLR8 (mTLR8) has long been considered unable to detect nucleic acids. Here, we show that mTLR8 is an active receptor that responds to the synthetic agonist TL8-506 and the endogenous cyclic nucleotide 2′,3′-cGMP, with signalling potentiated by cooperative binding of nucleic acid fragments at the secondary ligand-binding site. Using *Tlr7*-deficient mice expressing *mTlr8* under control of the *Tlr7* promoter, we demonstrate that mTLR8 partially restores TLR7-dependent tonic transcriptional programs in splenocytes, indicating responsiveness to physiological ligands *in vivo*. Comparison with a chimeric human-mouse TLR8 knock-in model expressing human TLR8 from the endogenous murine *Tlr7* locus demonstrated that the mTLR8 ectodomain is intrinsically less sensitive, consistent with sequence divergence in the LRR11-12 region that links agonist recognition to antagonist regulation. Collectively, these findings establish mTLR8 as a functional nucleic acid sensor specialized for combinatorial detection of nucleic acid degradation products, extending the murine endosomal nucleic acid-sensing repertoire beyond TLR3, TLR7, TLR9, and TLR13.

## Introduction

The innate immune system utilizes specialized sensors, such as Toll-like receptors (TLRs), to detect specific molecular features and initiate a rapid response to pathogens^1,2^. TLR activation leads to the induction of downstream signaling pathways, which in turn result in the production of cytokines and other pro- and anti-inflammatory molecules. These processes drive immediate host defensive responses such as inflammation, orchestrate antigen-specific adaptive immune responses, and modulate both tissue injury and repair^3,4^. Thirteen TLRs are known in mammals, five of which - TLR3, 7, 8, 9, and 13 - are implicated in the detection of endosomal nucleic acids^2,5^. TLR3 senses viral and synthetic double- stranded RNAs (dsRNA), while TLR7 and human TLR8 detect RNA degradation fragments from pathogens and endogenous single-stranded RNA (ssRNA), including siRNA and microRNA (miRNA), and synthetic base analogs^6–10^. TLR9 recognizes CpG-rich single-stranded DNA (ssDNA), while TLR13, exclusively expressed in rodents, binds specific sequences within bacterial ribosomal RNA^11,12^.

Thought to be the result of a duplication on chromosome X (ChX)^13^, TLR7 and TLR8 share a common mode of activation by RNA degradation products binding to two sites. Site 1 is highly conserved between TLR7 and TLR8 and binds certain nucleosides including guanosine (G, TLR7) and its derivative 2′,3′-cyclic guanosine monophosphate (2′,3′-cGMP, TLR7) and uridine (U, TLR8), as well as base analogs, such as loxoribine (TLR7), CL075, R848 (TLR7/TLR8), and TL8-506 (murine (m)TLR7/human (h)TLR8). Site 2 binds short U-rich motifs (e.g., UUU [TLR7], UG [TLR8]) and DNA fragments (TLR8) which cooperate with site 1 to enhance receptor activation^14–20^. Cooperation of site 1 and 2 engagement by nucleic acid fragments is necessary to outcompete basal antagonism of these receptors, enacted by binding of 2’-O-methyl guanosine RNA fragments, that safeguards against autoimmunity and autoinflammation^19,20^.

Although human and mouse TLR8 share high sequence homology, mTLR8 has long been considered unable to detect nucleic acids. Early functional characterization showed that TLR7-deficient mice fail to respond to canonical hTLR8 agonists, such as U, ssRNA40, or the synthetic imidazoquinoline R848^10,21–23^. However, these early studies predated our current structural understanding of hTLR8’s cooperative site 1 and site 2 activation. As rodents additionally express TLR13, which binds to bacterial RNA ligands otherwise sensed by hTLR8, it was concluded that mice can spare TLR8 function^24^. Nonetheless, several studies published as early as 2006 pointed to a role of functional mTLR8 in the context of brain development, neuronal injury, neuropathic pain, and autoinflammation^25–27^. Notably, *Tlr8*-deficiency was found to result in increased *Tlr7* function promoting autoimmunity, indicating that its presence in dendritic cells is necessary to restrict TLR7 function^26^, presumably through limiting TLR7 trafficking by UNC93B1^26,28,29^. Potentiation of hTLR8 site 1 synthetic agonists by site 2 ligands, such as thymidine-rich DNA (poly-dT), is conserved in mTLR8^23,30,31^, suggesting that despite differences between these orthologs, mTLR8 can also be activated through cooperative binding of nucleic acid fragments.

Here, we comprehensively analyze site 1 and site 2 cooperative sensing in mTLR8 and make use of a novel mouse model placing mTLR8 under the control of the *Tlr7* promoter. Our data demonstrate that, as with hTLR7, hTLR8, and mTLR7, mTLR8 can function as a sensor of nucleic acid fragments, including the site 1 agonist 2′,3′-cGMP, and exhibits activity partially overlapping with that of mTLR7.

## Results

### TL8-506 is a potent and direct mTLR8 agonist

The combination of poly-dT and the thiazoloquinolone derivative CL075, a dual TLR7/8 agonist^32,33^, has previously been shown to activate mTLR8 and suppress mTLR7 activity^31^, akin to what is seen with hTLR7 and hTLR8^20,34^. Here, we systematically analyzed modes of mTLR8 activation, employing commercial HEK293 cells stably co-expressing mTLR8 (or mTLR7, hTLR8, hTLR7 in later approaches) and an NF-κB/AP1-inducible secreted embryonic alkaline phosphatase (SEAP) reporter gene. Co- stimulation with several synthetic compounds previously identified as site 1 agonists for m/hTLR7 and/or hTLR8^15–18^, with or without poly-dT (Fig. 1A; Fig. S1A), revealed that the benzo-azepine analog TL8- 506^35^, which is slightly structurally larger than R848 (Fig. 1A), uniquely activated mTLR8 on its own. In agreement with prior reports, all synthetic agonists tested here activated mTLR8 sensing when combined with poly-dT^23,30,31^, except for loxoribine (Fig. 1A). Conversely, all the synthetic agonists tested activated HEK mTLR7 cells on their own, and this was significantly inhibited by poly-dT for TL8-506, Selgantolimod, imiquimod, and CL075 (Fig. 1B). Activation of HEK mTLR8 cells by TL8-506 was achieved at concentrations ≥500 ng/ml but were reduced to as low as 10 ng/ml when combined with poly-dT (Fig. 1C). Poly-dT potentiation of TL8-506 sensing also resulted in a faster mTLR8 response compared to TL8-506 alone (Fig. S1B). Conversely, poly-dT stably suppressed activation of HEK mTLR7 cells by TL8-506 over time (Fig. S1C, S1D), aligning with its direct antagonistic effect on hTLR7^20^. TL8-506 did not activate NF-κB signaling in the absence of mTLR7 or mTLR8 expression in the parental lines *Null2-k* and *Null1-v* (Fig. S1D, S1E).

**Figure 1.**
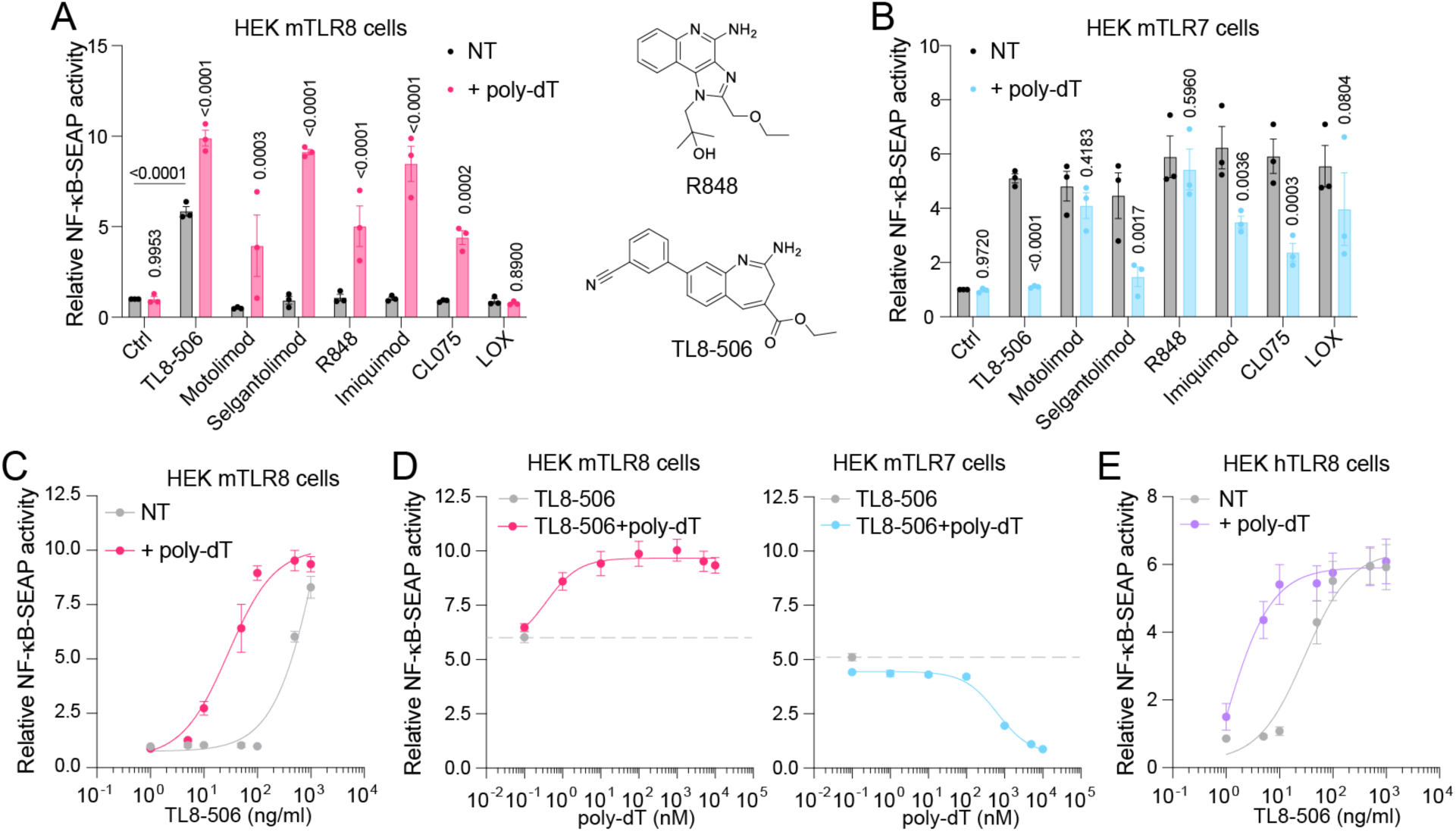
Combination of TL8-506 and poly-dT activates mTLR8 and hTLR8, while suppressing mTLR7 activity. (**A-B**) HEK-Blue cells co-expressing murine TLR8 (mTLR8) (**A**) or murine TLR7 (mTLR7) (**B**) and an NF-κB/AP1-inducible secreted embryonic alkaline phosphatase (SEAP) reporter gene were incubated overnight with TL8-506 (500 ng/ml), Motolimod (1 µM), Selgantolimod (1 µM), R848 (10 µg/ml), Imiquimod (5 µg/ml), CL075 (4 µM), or loxoribine (LOX; 1 mM), alone or in combination with deoxythymidine (poly-dT) 17-mer (5 µM). (**A, right**) Chemical structures of R848 and TL8-506 are shown. (**C**) mTLR8 reporter cells were incubated with various concentrations of TL8-506 alone or in combination with poly-dT 17-mer (5 µM) as indicated, for 24 h. (**D**) mTLR8 reporter cells (left) or mTLR7 reporter cells (right) were stimulated with TL8-506 (500 ng/ml) alone or in combination with various 17- mer poly-dT concentrations, as indicated, for 24 h (**D**). (**E**) HEK human TLR8 (hTLR8) reporter cells were incubated with various concentrations of TL8-506, alone or in combination with poly-dT (5 µM), as indicated, for 24 h. (**A**-**E**) Optical density was assessed the next day. Unstimulated cells (Ctrl) served as negative control. Data are mean of *n* = 3 (A, B, D right) or *n* = 4 (C, D left, E) independent experiments. Data are shown as fold change of the optical density normalized to unstimulated cells (Ctrl) (± s.e.m. and [**A-B**] two-way ANOVA with uncorrected Fisher’s LSD tests shown comparing conditions of treatments with or without poly-dT or otherwise indicated pairs of conditions).

Activation of mTLR8 by TL8-506 was robustly enhanced starting from 1 nM poly-dT, while mTLR7 suppression was observed at ≥1 μM poly-dT (Fig. 1D). Notably, in these cell models, HEK cells expressing hTLR8 could respond to lower concentrations of TL8-506 than those expressing mTLR8, as downstream signaling was detectable from ≥50 ng/ml (Fig. 1E). Similar hTLR8 responses were observed with ∼10 times less TL8-506 when combined with poly-dT, confirming that cooperative sensing increased sensitivity for both human and mouse receptors to a similar degree, noting that poly-dT alone had no effect on either receptor (Fig. S1A, S1F). Collectively, these results demonstrated that unlike other small molecule agonists of mTLR7 and hTLR8, TL8-506 can activate mTLR8 without cooperative engagement of poly-dT. This suggests that mTLR8 site 1 can function independently, albeit with distinctively reduced activity compared to its human counterpart.

### 2′,3′-cGMP and 2′,3′-cUMP are preferred natural agonists of mTLR8 site 1

The ability of TL8-506 to activate mTLR8 independently led us to examine whether natural nucleosides and their derivatives could function as endogenous mTLR8 agonists, given that G and 2′,3′-cGMP activate mTLR7 and hTLR7, whereas U and the pathogen-derived methylthioinosine (MTI) activate hTLR8 via site 1 binding^14,36^. We tested panels of RNA nucleosides (Fig. 2A), modified RNA nucleosides (Fig. 2B), 2′,3′-cyclic nucleotide monophosphates (cNMP) naturally produced upon endosomal enzymatic cleavage of RNAs^37^, and 3’,5’-cNMPs (Fig. 2C, 2D), with or without poly-dT, in HEK mTLR8 reporter cells. While U, pseudo-uridine (Ψ), 5-Methyluridine (m^5^U), 2′,3′-cUMP, and 2′,3′-cGMP all significantly activated mTLR8 in the presence of poly-dT, only 2′,3′-cGMP significantly induced mTLR8 activation in the absence of poly-dT (Fig. 2A-C). Co-stimulation with poly-dT and either 2′,3′-cUMP or 2′,3′-cGMP produced the strongest mTLR8-dependent responses, exceeding those induced by U by more than two-fold, with significant cooperative activation observed from 1 mM 2′,3′-cGMP (Fig. 2C; Fig. S2A). Poly-dT enhanced the response of HEK mTLR8 cells to 2′,3′-cGMP by >4-fold at concentrations as low as 3 mM, whereas 10 mM U was required to achieve a comparable effect (Fig. 2E; Fig. S2B). Despite this enhanced cooperativity relative to U, mTLR8 activation by 2′,3′-cGMP or 2′,3′-cUMP remained less potent than hTLR8 activation by U, for which poly-dT-mediated potentiation was detectable at concentrations as low as 0.1 mM U (Fig. S2C). Analysis of related nucleotide derivatives, including 3′-GMP, 2′-GMP, 5′-GMP, 3′-UMP, and 2′-UMP, demonstrated that only 2′,3′-cGMP and 2′,3′- cUMP elicited significant mTLR8 activation with poly-dT (Fig. 2F).

**Figure 2.**
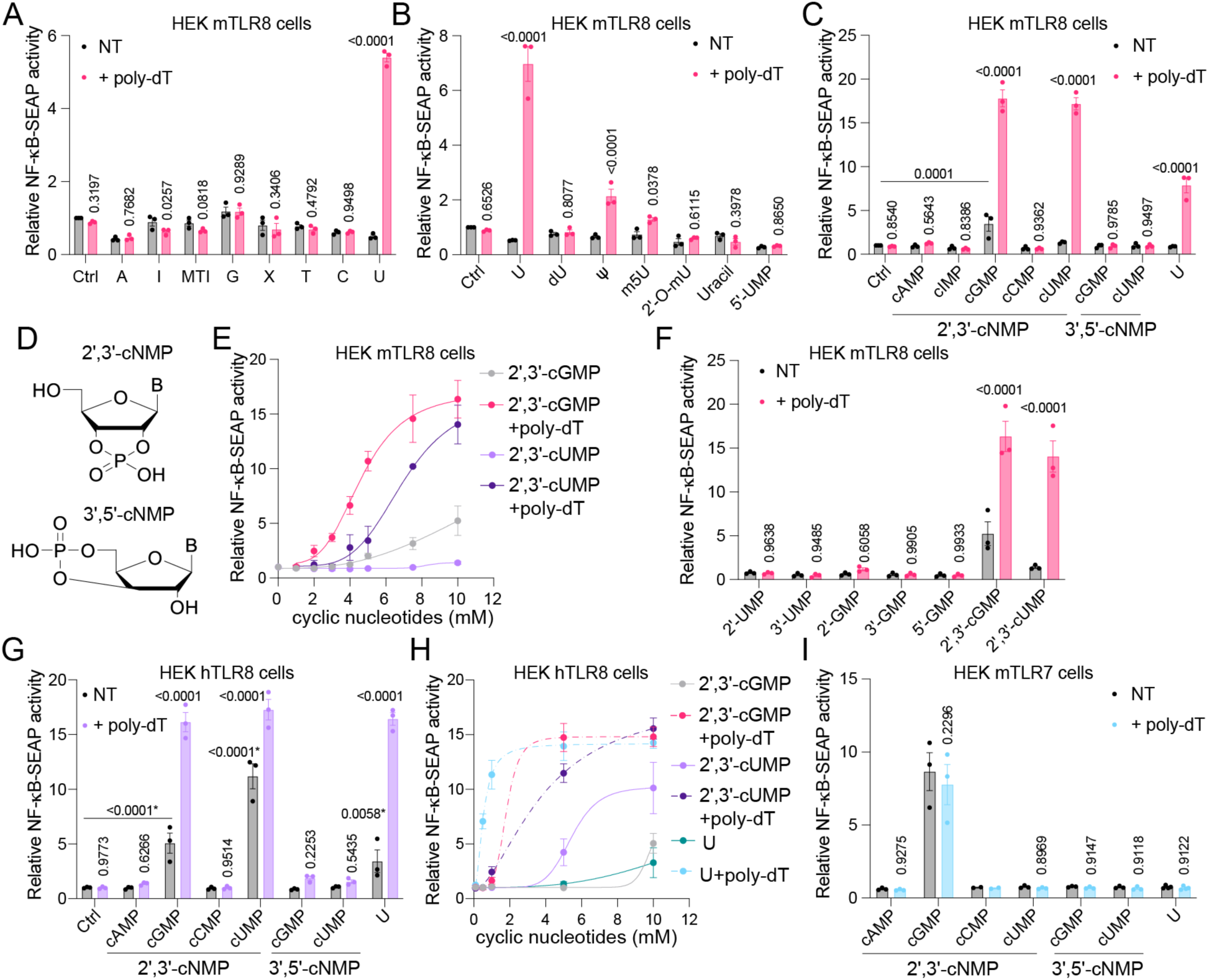
mTLR8 response to uridine, 2′,3′-cGMP, and 2′,3′-cUMP. (**A-C**) HEK mTLR8 reporter cells were incubated with (**A**) adenosine (A; 30 mM), inosine (I; 3.3 mM), methylthioinosine (MTI; 3.3 mM), guanosine (G; 1 mM), xanthosine (X; 1 mM), thymidine (T; 30 mM), cytidine (C; 30 mM), or uridine (U; 30 mM), alone or in combination with 17-mer poly-dT (5 µM), (**B**) U (30 mM), deoxyuridine (dU; 30 mM), pseudouridine (Ψ; 30 mM), 5’-methyluridine (m5U; 30 mM), 2’-O-methyluridine (2’-O-mU; 30 mM), uracil (30 mM), or U monophosphate (5’-UMP; 30 mM), alone or in combination with 17-mer poly-dT (5 µM), or (**C**) 2′,3′-cyclic nucleotide monophosphates (2′,3′-cNMPs, 10 mM) of A (2′,3′-cAMP), I (2′,3′-cIMP; 5 mM), G (2′,3′-cGMP), C (2′,3′-cCMP), U (2′,3′-cUMP), non-cyclic U (10 mM), 3’,5’-cNMPs (10 mM) of G (3’,5’-cGMP), or U (3’,5’-cUMP), alone or in combination with 17-mer poly-dT (5 µM), for 24 h. (**D**) Structure of 2′,3′-cNMP and 3’,5’-cNMP. (**E-F**) HEK mTLR8 reporter cells were incubated with (**E**) various concentrations of 2′,3′-cGMP or 2′,3′-cUMP, as indicated, alone or in combination with 17-mer poly-dT (5 µM), or (**F**) with the indicated nucleotides and their derivatives, including 3′-GMP, 2′-GMP, 5′- GMP, 3′-UMP, and 2′-UMP (at 10 mM each), alone or in combination with 17-mer poly-dT (5 µM), for 24 h. (**G-H**) hTLR8 reporter cells were incubated for 24 h with (**G**) 2′,3′-cAMP, 2′,3′-cCMP, 2′,3′-cGMP, 2′,3′- cUMP, 3’,5’-cGMP, 3’,5’-cUMP, or U, alone (all 10 mM) or in combination with 17-mer poly-dT (5 µM), or (**H**) with various concentrations of 2′,3′-cGMP, 2′,3′-cUMP, or U, as indicated, alone or in combination with 17-mer poly-dT (5 µM). (**I**) HEK murine TLR7 (mTLR7) reporter cells were incubated with 2′,3′-cyclic nucleotide monophosphates (2′,3′-cNMPs, 10 mM) of A (2′,3′-cAMP), G (2′,3′-cGMP), C (2′,3′-cCMP), U (2′,3′-cUMP), 3’,5’-cNMPs (10 mM) of G (3’,5’-cGMP), or U (3’,5’-cUMP), or non-cyclic U (10 mM), alone or in combination with 17-mer poly-dT (5 µM), for 24 h. (**A-I**) Optical density was assessed the next day. Unstimulated cells (Ctrl) served as negative control. Data are mean of *n* = 3 independent experiments. Data are shown as fold change of the optical density normalized to unstimulated cells (Ctrl) (± s.e.m. and [A-C, F-G, I] two-way ANOVA with uncorrected Fisher’s LSD tests shown comparing conditions of treatments with or without poly-dT or otherwise indicated pairs of conditions; for G, *P* values with a * indicate comparisons to Ctrl condition).

Notably, unlike mTLR8, hTLR8 responded to 2′,3′-cUMP and U on their own, and similarly to mTLR8, it responded to 2′,3′-cGMP (Fig. 2G; Fig. S2C). While 2′,3′-cUMP was the strongest hTLR8 agonist on its own, cooperative stimulation with poly-dT significantly enhanced the hTLR8 response to U, 2′,3′-cGMP and 2′,3′-cUMP (Fig. 2H). With respect to mTLR7, 2′,3′-cGMP was the only 2′,3′-cNMP tested able to activate the receptor at the concentration tested, and the response was not enhanced by poly-dT (Fig. 2I). Collectively, these findings indicate that mTLR8 is less sensitive than hTLR8 to site 1 ligands U and 2′,3′-cUMP, whereas both receptors respond to 2′,3′-cGMP. Notably, cooperative sensing with poly-dT is conserved across species and increases sensitivity of mTLR8 to U, 2′,3′-cUMP, or 2′,3′- cGMP.

## Site 2-mediated potentiation of mTLR8 sensing by DNA and RNA fragments

We next focused on characterization of site 2 ligands, beyond the case exemplified above for poly-dT. Similar to what has been reported for hTLR8^20,30,34^, mTLR8 sensing of low-dose TL8-506 was significantly increased with poly-dT oligomers ≥9 bases (Fig. 3A). Beyond poly-dT, poly-dA induced comparatively mild activation, while poly-dC and poly-dG failed to induce significant receptor activation, and none of the tested oligonucleotides induced a mTLR8 response on their own (Fig. S3A). A low but significant activation of mTLR8 was also observed with poly-U, but not poly-dU, in the presence of TL8- 506 (FC ∼2), while no activation was observed on their own (Fig. S3B). Conversely, strong potentiation of TL8-506 sensing was seen with poly-U and poly-dU on hTLR8 (Fig. S3C). To confirm these results and gain further mechanistic understanding of mTLR8 potentiation, we also analyzed the activity of a 3- base oligonucleotide, _m_G_d_C_d_C, which we recently established as a direct binder of hTLR8 site 2 through crystallographic analyses^19^. Unlike hTLR8, mTLR8 sensing of TL8-506 was not potentiated by _m_G_d_C_d_C; however, elongation of the 3-mer to ≥6-bases with deoxycytidine moieties (e.g. _m_G_d_C_d_C_c_C_c_C_c_C) was sufficient to result in significant potentiation of TL8-506 sensing by mTLR8 (Fig. 3B), indicating close structural similarities in site 2 between human and mouse TLR8.

**Figure 3.**
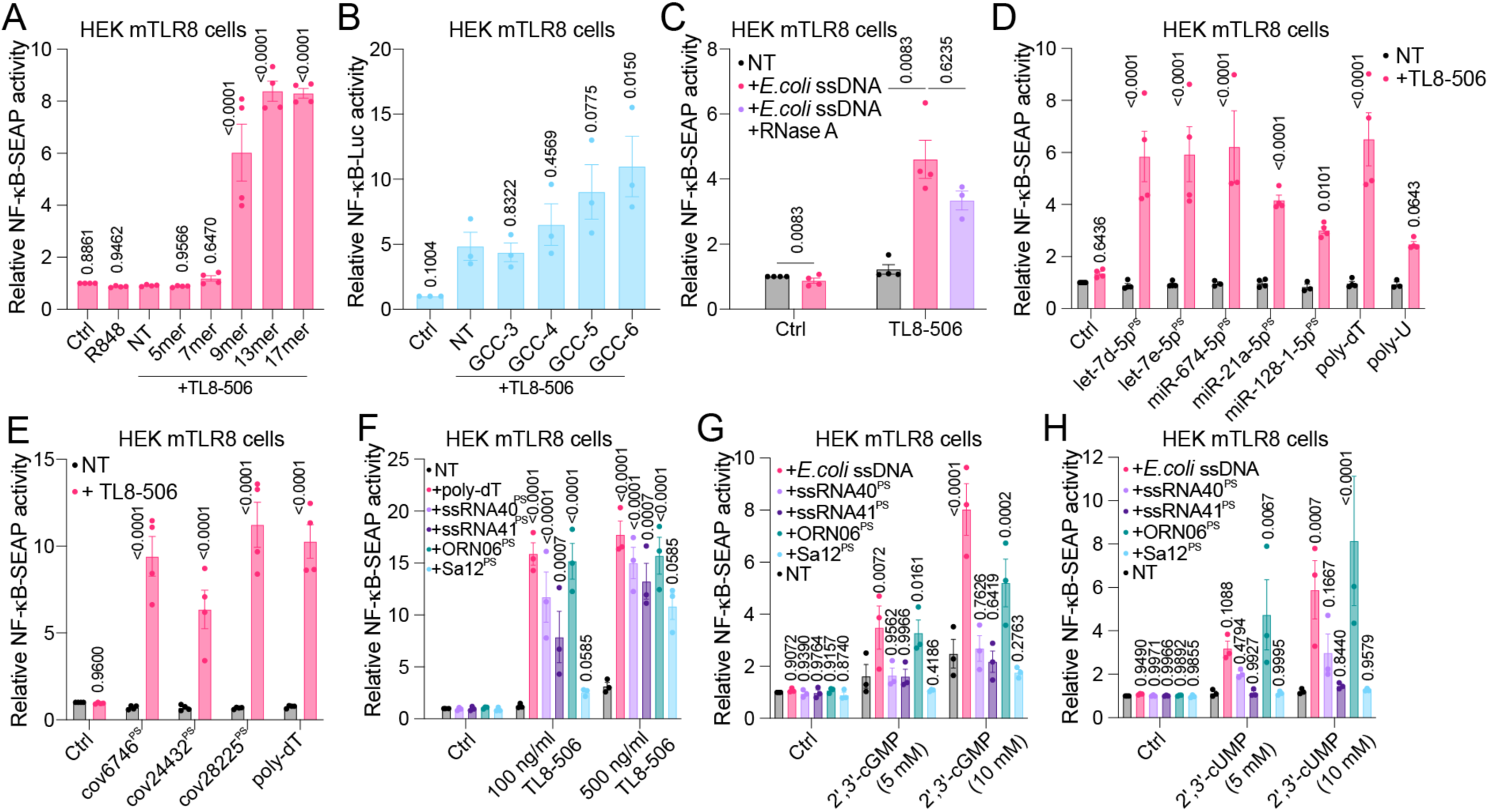
Sequence-dependent potentiation of mTLR8 sensing by short ssDNAs and ssRNAs. HEK mTLR8 reporter cells were incubated overnight with (**A**) TL8-506 (100 ng/ml), alone or in combination with different poly-dT oligonucleotides with increasing lengths in thymidine (5 µM), as indicated, or R848 (5 µg/ml); (**B**) TL8-506 (100 ng/ml), alone or in combination with 3-base oligonucleotide _m_G_d_C_d_C^PS^ (GCC-3) or 4, 5- and 6-mer _m_G_d_C_d_C derivatives (5 µM); (**C**) *E. coli* ssDNA (20 µg/ml) in combination with TL8-506 (100 ng/ml), or TL8-506 plus RNase A (10 µg/ml); (**D, E**) Transfected miRNAs (**D**) or SARS-CoV-2 fragments (**E**, 10 µg/ml), or 17-mer poly-dT and poly-U (5 µM), alone or in combination with TL8-506 (100 ng/ml); (**F-H**) poly-dT (5 µM) or *E. coli* ssDNA (10 µg/ml), ssRNA40^PS^ (10 µg/ml), ssRNA41^PS^ (10 µg/ml), ORN06^PS^ (10 µg/ml), or Sa12^PS^ (10 µg/ml), alone or in combination with 100 ng/ml TL8-506 or 500 ng/ml TL8-506, as indicated (**F**), or in combination with 5 mM or 10 mM 2′,3′-cGMP (**G**) or 2′,3′-cUMP (**H**), as indicated. (**A-H**) NT is non-treated. (**A-G**) Optical density was assessed the next day. Data are mean of *n = 3* (B, F-H) or *n = 4* (A, C-E) independent experiments. Data are shown as fold change of the optical density normalized to unstimulated cells (Ctrl) (± s.e.m. and [A-B] one-way or [C-H] two-way ANOVA with uncorrected Fisher’s LSD tests shown comparing conditions to TL8-506 only condition [A-B, F], comparing treatments with or without TL8-506 [D-E], comparing treatments to NT conditions [G, H] or otherwise indicated pairs of conditions [C]).

We next investigated whether mTLR8 sensing could also be potentiated by purified bacterial single-stranded (ss)DNA, an established TLR9 ligand. Co-incubation of ssDNA fragments purified from *E. coli* with TL8-506 significantly potentiated mTLR8 activation in a dose-dependent manner, while *E. coli* ssDNA alone failed to activate the receptor (Fig. 3C; Fig. S3D). RNase A treatment of the ssDNA did not significantly affect this potentiation, suggesting this effect was independent of RNA contamination (Fig. 3C). Importantly, unlike its murine counterpart, hTLR8 sensing of TL8-506 was not potentiated further by *E. coli* ssDNA at the dose used (Fig. S3E).

As ssRNA degradation products provide ligands for both ligand-binding sites of hTLR8 and contribute to TLR7 activation^15,18,37^, we next investigated whether mTLR8 could also recognize biologically relevant host- and pathogen-derived RNA sequences. These included the host-derived miRNAs let-7d-5p, let-7e-5p, miR-128-1-5p, miR-21a-5p, and miR-674-5p, as well as SARS-CoV-2- derived RNAs (cov6748, cov24432, and cov28225), previously implicated in hTLR8 and/or TLR7 activation^8,38,39^. All tested phosphorothioate (PS)-modified oligoribonucleotides potentiated TL8-506- induced mTLR8 activation, but did not activate mTLR8 on their own, suggesting engagement of the second ligand-binding site of mTLR8 (Figs. 3D, 3E). Next, we extended our analysis to established, sequence-defined hTLR8-potentiating ssRNAs, including ssRNA40^PS^ and ORN06^PS^, as well as the uridine-free ligands ssRNA41^PS^ and Sa12^PS^ ^14^. While none of these PS-modified RNAs activated mTLR8 on their own, ORN06^PS^ strongly potentiated receptor activation at low TL8-506 concentrations, followed by ssRNA40^PS^ and, to a lesser extent, ssRNA41^PS^, whereas Sa12^PS^ showed no detectable effect. At higher TL8-506 concentrations, however, all four ligands enhanced mTLR8 activation (Fig. 3F). These responses largely mirrored the ligand preferences reported for hTLR8^14^, suggesting similar sequence- dependent recognition at site 2^18^. Finally, to determine whether site 2-mediated potentiation also extends to naturally occurring site 1 agonists, we analyzed the combined effects of *E. coli* ssDNA and PS- modified ssRNA40^PS^, ssRNA41^PS^, ORN06^PS^, or Sa12^PS^, with 2′,3′-cGMP and 2′,3′-cUMP. Among these, *E. coli* ssDNA and ORN06^PS^ were the only nucleic acids to significantly potentiate mTLR8 activation by both cyclic nucleotides in a dose-dependent manner (Figs. 3G, 3H).

Collectively, these findings demonstrated that mTLR8 responds to a broad range of DNA and RNA fragments through cooperative ligand recognition, with sequence-dependent preferences largely resembling those of hTLR8. Moreover, this potentiation is not restricted to synthetic agonists such as TL8-506 but also extends to naturally occurring cyclic nucleotides, supporting a shared mechanism of cooperative site 1 and site 2 ligand recognition.

## mTLR8 expressed in murine BMDMs responds to nucleic acid fragments

Similar to hTLR8, mTLR8 is predominantly expressed in the myeloid compartment^40^. We therefore explored how mTLR8 might respond to nucleic acid fragments in murine bone marrow-derived macrophages (BMDMs). TL8-506 alone failed to induce a response at concentrations below 500 ng/mL but synergized with poly-dT at low doses to trigger TNF production in murine wild-type (WT) BMDMs, suggesting mTLR8 activity in these cells (Fig. 4A). This response was abolished in *Tlr7/8* double- deficient (Tlr7/8 DKO) BMDMs (Fig. 4B). In *Tlr7*^⁻/⁻^ (7KO) BMDMs (mTLR8-competent), TNF production occurred only with co-stimulation by TL8-506 and poly-dT at the doses used (Fig. 4C). Conversely, *Tlr8*^⁻/⁻^ (8KO) BMDMs (TLR7-competent) responded to TL8-506 alone, but this response was blunted by poly- dT (Fig. 4D). Furthermore, while 2′,3′-cGMP alone induced a strong response from WT BMDMs and 8KO cells that was inhibited by poly-dT (Figs. 4E, 4F), 7KO cells showed a weak response to 2′,3′-cGMP that was enhanced by poly-dT (Fig. 4G). Tlr7/8 DKO BMDMs were not responsive to 2′,3′-cGMP alone or in combination with poly-dT or ssRNA40^PS^ (Fig. 4H). Notably, 8KO BMDMs mirrored the response pattern of WT cells, implying that TLR7 is the dominant sensor of 2′,3′-cGMP in BMDMs (Figs. 4E, 4F). Importantly, poly-dT significantly sensitized WT and 7KO BMDMs to 2′,3′-cUMP stimulation that was absent in 8KO cells, suggesting that site 2 binders may also be recruited to mTLR8 for activation by 2′,3′-cUMP in BMDMs (Fig. 4I). In 7KO BMDMs, ORN06^PS^ and ssRNA40^PS^ potentiated TL8-506-induced mTLR8 responses like poly-dT but did not induce any response on their own (Fig. 4J). Conversely, unlike poly-dT, ORN06^PS^ and ssRNA40^PS^ strongly activated 8KO and WT BMDMs in the absence of TL8-506, and there was no robust further potentiation of TNF production in combination with TL8-506 with these RNAs (Fig. 4J; Fig. S4A). *E. coli* ssDNA enhanced TL8-506 and 2′,3′-cGMP sensing in 7KO and *Tlr3/7/9^⁻/⁻^* (3/7/9KO) BMDMs (Figs. 4K, 4L), consistent with TLR9-independent bacterial ssDNA detection by mTLR8.

**Figure 4.**
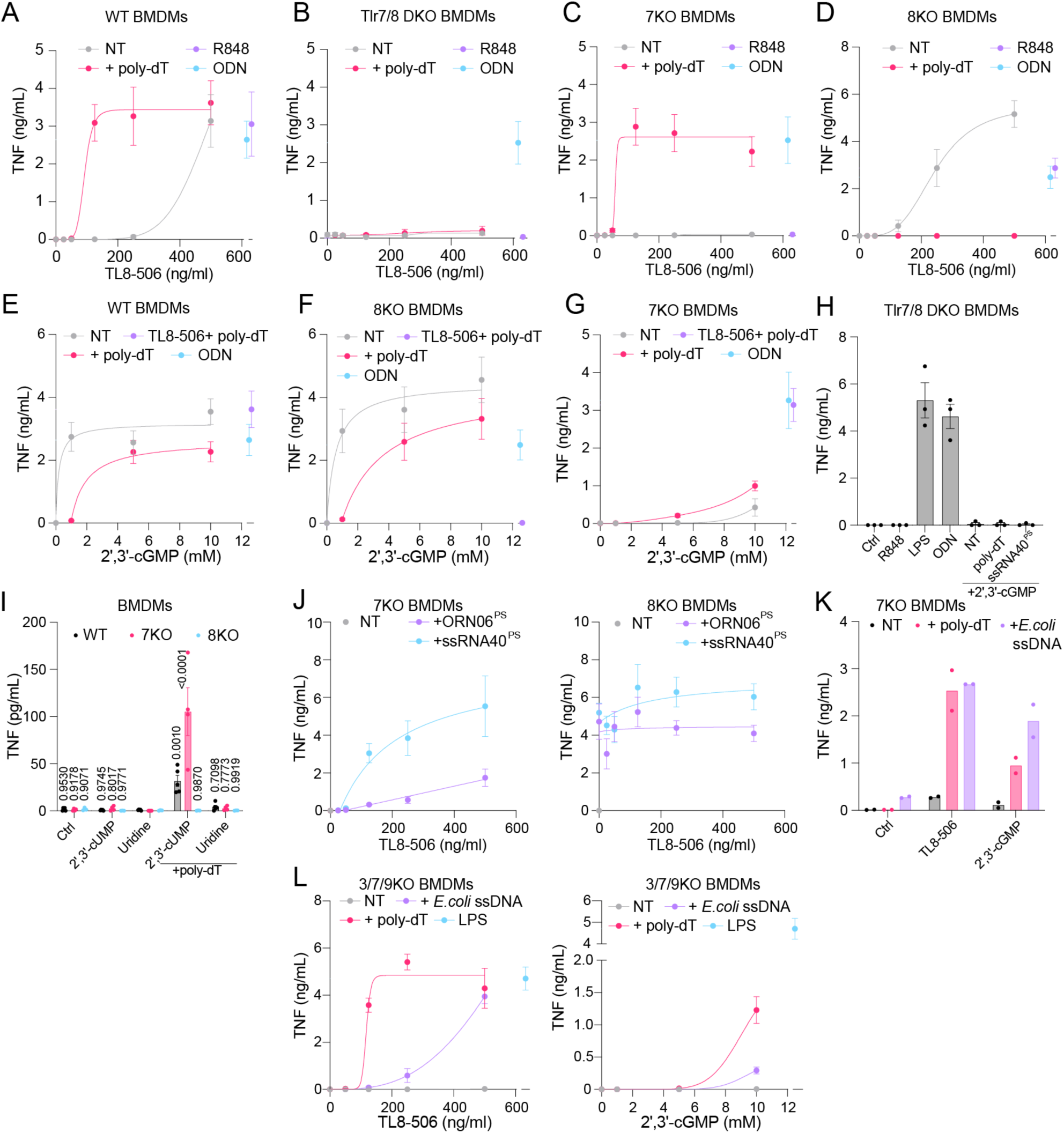
Murine bone marrow-derived macrophages respond to 2′,3′-cGMP and to combinations of cyclic nucleotides, nucleosides, ssRNA, and ssDNA through TLR8. Bone marrow-derived macrophages (BMDMs) isolated from (**A**) C57BL/6 (wild-type, WT), (**B**) Tlr7/8 DKO, (**C**) *Tlr7*^⁻/⁻^ (7KO), or (**D**) *Tlr8*^⁻/⁻^ (8KO) mice were incubated overnight with various concentrations of TL8-506, as indicated, with or without 20-mer poly-dT (2.5 µM), R848 (10 µg/ml), or ODN 1826 (500 nM). BMDMs from (**E**) WT, (**F**) 8KO, or (**G**) 7KO mice were incubated with indicated dose of 2′,3′-cGMP alone or in combination with 20-mer poly-dT (2.5 µM), with TL8-506 (500 ng/ml) + 20-mer poly-dT (2.5 µM), or ODN 1826 (500 nM) for 16 h. (**H**) Tlr7/8 DKO BMDMs were incubated overnight with 10 mM 2′,3′-cGMP alone or in combination with 20-mer poly-dT (3 µM) or transfected with ssRNA40^PS^ (10 µg/ml), R848 (1 µg/ml), LPS (100 ng/ml), or ODN 1826 (500 nM). (**I**) BMDMs from WT, 7KO, or 8KO mice, as indicated, were incubated overnight with 2′,3′-cUMP (10 mM), uridine (10 mM), alone or in combination with 20-mer poly-dT (2.5 µM). (**J**) BMDMs from 7KO (left) and 8KO (right) mice were incubated overnight with various TL8-506 concentrations, alone or in combination with 20-mer poly-dT (2.5 µM), transfected ORN06^PS^ (10 µg/ml) or ssRNA40^PS^ (10 µg/ml), as indicated. (**K**) 7KO BMDMs were incubated overnight with TL8- 506 (500 ng/ml) or 2′,3′-cGMP (10 mM), alone or in combination with 20-mer poly-dT (2.5 µM) or *E. coli* ssDNA (10 µg/ml). (**L**) TLR3/7/9 TKO (3/7/9KO) BMDMs were incubated overnight with various doses of TL8-506 (left) or 2′,3′-cGMP (right), as indicated, alone or in combination with 20-mer poly-dT (2.5 µM) or *E. coli* ssDNA (10 μg/ml) for 16 h. LPS (100 ng/ml) served as positive control. (**A-L**) TNF concentration levels in cell supernatants were assessed by ELISA. Unstimulated cells (NT) served as negative control. Data are mean of *n* = 2 (**K**), *n =* 3 (B, D, F, H, J), *n =* 4 (A, E, G, I, L), or *n =* 5 (C) independent experiments (± s.e.m.).

To extend our analyses beyond TNF responses, we compared 15 cytokines produced by WT, 7KO, and 8KO BMDMs stimulated with TL8-506 or 2′,3′-cGMP in the presence or absence of poly-dT (Fig. S4B). Except for VEGF-A, all cytokines were induced by TL8-506 or 2′,3′-cGMP in WT and 8KO cells, whereas responses were reduced in 7KO cells, consistent with the dominant role of TLR7 in sensing these agonists at the dose used (Fig. S4B). Poly-dT co-stimulation enhanced cytokine production in 7KO cells in response to both agonists but reduced TL8-506-induced responses in WT and 8KO BMDMs, with a stronger inhibitory effect in 8KO cells, consistent with the antagonistic effect of poly-dT on TLR7 signaling (Fig. S4B).

To determine whether activation of mTLR8 can be induced by the mTLR8 agonists identified in this study *in vivo*, we next utilized an established brain injection model in which receptor activation is assessed following administration of the respective TLR agonist^39,41–43^. To this end, WT, 8KO, and 7KO mice were intrathecally injected with the combination TL8-506 plus poly-dT or 2′,3′-cGMP (Figs. S4C, S4D). Three days after injection, immunohistochemical analysis of the cerebral cortex revealed a significant increase in microglial cells (Iba1^+^, Fig. S4C), the major immune cells in the brain, and loss of neurons (NeuN^+^, Fig. S4D) in WT mice injected with TL8-506 plus poly-dT or 2′,3′-cGMP compared to vehicle control condition. In 8KO mice, these effects induced by poly-dT plus TL8-506 were completely abolished. In contrast, injection of 2′,3′-cGMP into 8KO mice led to similar effects as observed in WT mice, although to a lesser extent. Conversely, in 7KO mice, 2′,3′-cGMP injection had no significant effect on microglial and neuronal numbers in the cortex, while the combination of TL8-506 and poly-dT induced significant microglial accumulation (Fig. S4C) and neuronal loss (Fig. S4D). These results align with our findings on mTLR7 as the dominant receptor for 2′,3′-cGMP signaling (see Figs. 2, 4E, 4F). Collectively, since microglial accumulation and concurrent reduction in neurons are hallmarks of neuroinflammation and neurodegeneration^44^, our findings support the selective pro-inflammatory and injurious function of mTLR8 in the brain, when engaged with its agonists such as the combination of poly-dT and TL8-506.

## mTLR8 can contribute to tonic basal nucleic acid sensing

Although mTLR7 is considerably more sensitive to 2′,3′-cGMP, DNA- or RNA-mediated site 2 engagement strongly enhanced mTLR8 sensing, prompting us to ask whether mTLR8 could compensate for the loss of mTLR7 since they can both detect 2′,3′-cGMP. Low-level tonic TLR7 signaling has been reported in select B-cell subsets^45,46^. To directly test whether mTLR8 could functionally replace TLR7, we generated a *Tlr7*^-/-^ knockout mouse model, in which *mTlr8* is expressed from the upstream adjacent *mTlr7* promoter, with the intent of concurrently increasing TLR8 expression in B cells and ablating TLR7 function (referred herein as the pTlr7-mTlr8 mouse, Fig. 5A). We complemented this model by generating a second chimeric strain in which the human TLR8 ectodomain was fused with the mTLR8 transmembrane and TIR domains and was similarly placed under *Tlr7* promoter control (referred to as the hTLR8KI mice hereafter), allowing head-to-head comparison of human and mouse TLR8 ectodomain function in primary cells (Fig. 5A). RT-qPCR analyses of BMDMs from these mice confirmed loss of *mTlr7* expression in pTlr7-mTlr8, hTLR8KI, and Tlr7/8 DKO mice (Fig. S5A). Notably, expressing *mTlr8* from the *mTlr7* promoter in pTlr7-mTlr8 mice only marginally increased *mTlr8* expression in these macrophages (Fig. S5A). Stimulation of BMDMs from pTlr7-mTlr8, hTLR8KI, Tlr7/8 DKO, and matched WT mice confirmed the function of the *hTLR8* fusion gene, which exhibited greatly increased sensitivity to TL8-506 compared to WT mice expressing endogenous mTLR7/mTLR8, as measured by TNF production (Fig. 5B). While TL8-506 failed to activate mTLR8 on its own at these doses, it synergised with poly-dT in pTlr7-mTlr8 mice, similarly to 7KO cells (Fig. 5C; see Fig. 4C). As shown in Fig. 5D, further functional validations of receptors in these mice were carried out confirming 1) loss of TLR7 function in pTlr7-Tlr8 and Tlr7/8 DKO mice, through loss of R848 sensing; 2) U sensing on its own and its potentiation by poly-dT or ssRNA40^PS^ in hTLR8KI mice; 3) ssRNA40^PS^ sensing on its own in WT (mTLR7) and hTLR8KI mice, and its potentiation by TL8-506 in pTlr7-mTlr8, WT, and hTLR8KI mice. We also confirmed 2′,3′-cGMP sensing of mTLR8 in combination with poly-dT or ssRNA40^PS^ in pTlr7-mTlr8 BMDMs (Fig. S5B).

**Figure 5.**
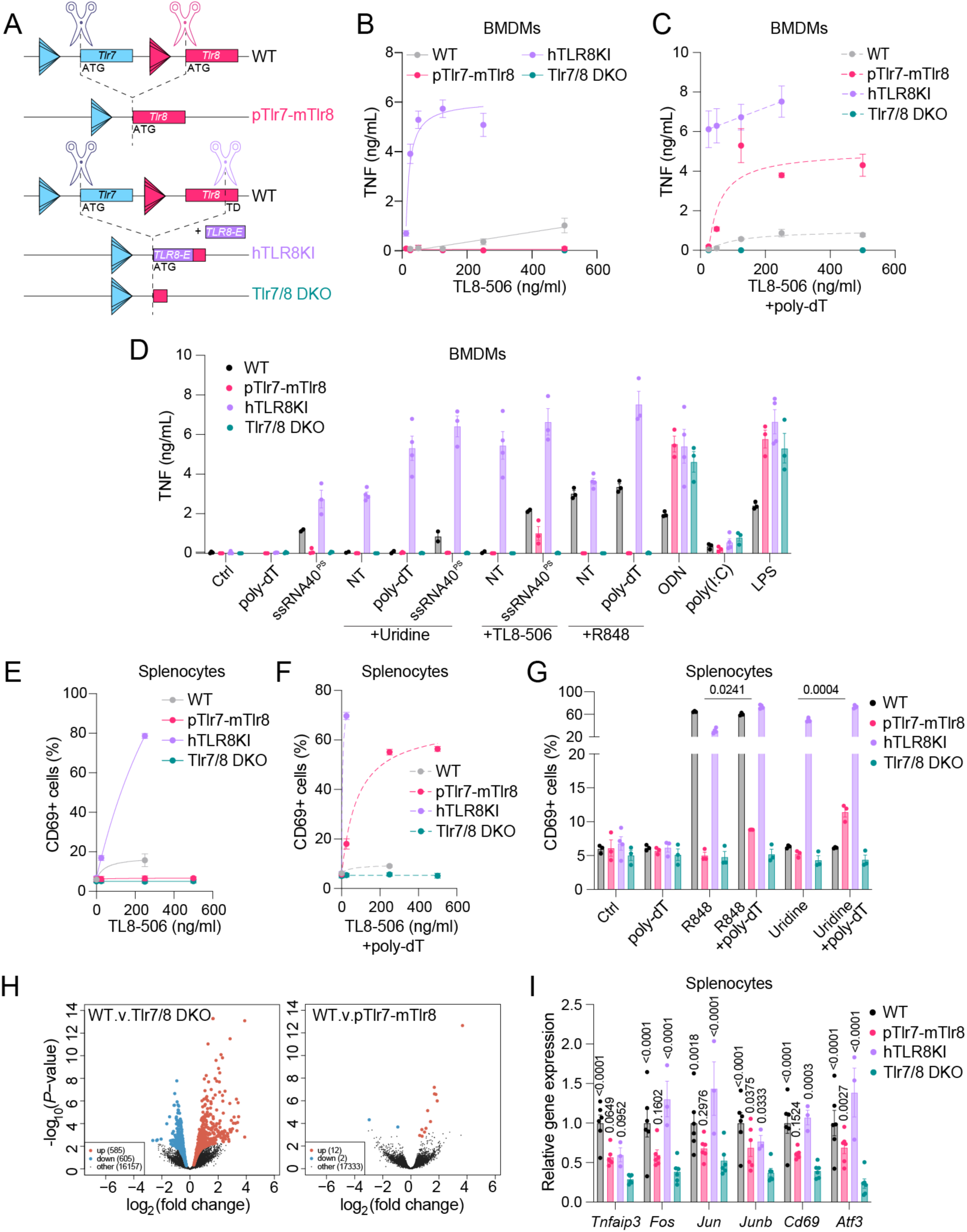
mTLR8 can contribute to tonic basal nucleic acid sensing. (**A**) CRISPR-Cas9 generation of pTlr7-mTlr8, hTLR8KI, and Tlr7/8 DKO mouse models. In pTlr7-mTlr8 mice, gRNAs targeting sequences flanking the first ATGs of *mTlr7* and *mTlr8* deleted the m*Tlr7* coding region and *mTlr8* promoter, placing *mTlr8* under *Tlr7* promoter control via repair oligo-mediated fusion. For hTLR8KI and Tlr7/8 DKO mice, gRNAs targeted the mTLR8 ectodomain-transmembrane domain (TD) (aa 827), and a synthetic hTLR8 ectodomain (*TLR8-E*) insert (aa 1-834) was introduced to generate a chimeric receptor containing murine transmembrane and TIR domains. Tlr7/8 DKO mice were generated by direct ligation without insertion of the *hTLR8* sequence. (**B, C**) BMDMs from 6-12-week-old hemizygous males of indicated genotype were pre-treated with (**C**) or without (**B**) 1 μM 20-mer poly-dT, or transfected with 750 nM ssRNA40^PS^ (**D**) for 30 min prior to overnight stimulation with indicated doses of TL8-506 (**B**, **C**), 20 mM U, 1 μg/mL R848, 500 nM ODN 1826 (ODN), 10 μg/mL poly (I:C) or 100 ng/mL LPS (**D**). TNF levels in cell supernatants were quantified by ELISA. Data are presented as mean ± s.e.m. of *n* = 3 (other genotypes) or 4 mice (hTLR8KI, **D**). (**D**) ssRNA40^PS^ conditions were conducted in *n* = 2 WT mice. (**E**) Primary splenocytes from 6-12-week-old hemizygous males of indicated genotype were untreated (**E**) or pre-treated with 1 μM 20-mer poly-dT for 30 min prior to stimulation with indicated doses of TL8- 506 (**E, F**), 1 μg/mL R848 or 20 mM U overnight. CD69^+^ cells were analysed by flow cytometer. Data are presented as mean ± s.e.m. of *n* = 3 (other genotypes) or 4 (hTLR8KI) mice. (**G**) Two-way ANOVA analyses with uncorrected Fisher’s LSD tests are shown. (**H**) Unstimulated splenocyte RNA from WT, Tlr7/8 DKO, and pTlr7-mTlr8 mice were analysed by RNA-sequencing. Volcano plot of the genes significantly different between Tlr7/8 DKO and WT cells (left), and pTlr7-mTlr8 and WT cells (right) are shown (blue are down-regulated and red are upregulated). (**I**) RT-qPCR analyses of *Tnfaip3, Fos, Jun, Junb, Cd69,* and *Atf3,* reported to *18S* in RNA lysates from primary splenocytes from 3 (hTLR8KI) or 6 (WT, pTlr7-mTlr8, or Tlr7/8 DKO) mice. Data are shown relative to WT mice (± s.e.m. and two-way ANOVA with uncorrected Fisher’s LSD tests shown compared to Tlr7/8 DKO mouse condition).

Consistent with these findings, splenocytes from hTLR8KI mice showed ∼10-fold greater sensitivity to TL8-506 than WT cells, as measured by the percentage of CD69⁺ cells (Fig. 5E). However, unlike BMDMs, which expressed similar levels of mTlr8 in pTlr7-mTlr8 and WT mice, splenocytes from pTlr7-mTlr8 mice expressed ∼3-fold higher *mTlr8* mRNA levels than WT splenocytes, supporting a *Tlr7* promoter-specific activity over that of *mTlr8* in the spleen (Fig. S5C). As a result, TL8-506 synergized strongly with poly-dT in pTlr7-mTlr8 but not in WT splenocytes, with mTLR8 activity detectable at concentrations as low as 25 ng/ml TL8-506 with poly-dT (Fig. 5F). Moreover, treatment with U plus poly- dT induced a modest but significant increase in CD69⁺ cells in pTlr7-mTlr8, not seen in WT, splenocytes (Fig. 5G). These findings demonstrate that mTLR8 becomes functionally responsive in splenocytes when expressed under the *Tlr7* promoter in pTlr7-mTlr8 mice.

To define whether this increased splenic mTlr8 expression could result in low-level tonic signalling, we next performed bulk RNA-seq of unstimulated splenocytes. Compared with WT cells, *Tlr7/8* deficiency caused widespread transcriptional changes (585 significantly upregulated and 605 downregulated genes), highlighting the importance of basal TLR7 signalling in the spleen (Fig. 5H; Fig. S5D). In contrast, transcriptomes of pTlr7-mTlr8 splenocytes closely resembled that of WT cells, differing in only 14 genes (12 upregulated, 2 downregulated) (Fig. 5H; Fig. S5D). RT-qPCR confirmed increase of several TNF-related genes (*Junb,* and *Atf3* being significant at *p < 0.05*) between pTlr7- mTlr8 and Tlr7/8 DKO splenocytes, indicating that mTLR8 partially compensates for the loss of TLR7 when expressed under the *Tlr7* promoter (Fig. 5I). This compensatory effect was even stronger in hTLR8KI splenocytes, consistent with the greater responsiveness of hTLR8 to RNA fragments compared to mTLR8 (Fig. 5I). Notably, although tonic mTLR8 signalling partially compensated for TLR7 deficiency at the transcriptional level in splenocytes, pTlr7-mTlr8 and Tlr7/8 DKO mice displayed comparable reductions in numbers of splenic germinal center B cells (GC B cells), plasmablasts, and atypical B cells (ABCs), without otherwise impacting spleen weight, paralleling prior findings in *Tlr7*- deficient mice^45,46^ (Figs. S5E, S5F).

Collectively, these genetic mouse models establish that mTLR8 can mediate both tonic and ligand-induced signalling independently of TLR7, although less efficiently than what is observed in our chimera mice expressing the hTLR8 ectodomain.

## mTLR8 lacks inhibition by established TLR7/8 antagonists

The reduced activity of mTLR8 has been attributed to the absence of the human-specific RQSYA motif^23^; however, our data show that mTLR8 retains responsiveness to the site 1 agonist 2′,3′-cGMP, a property shared with hTLR8. To define the molecular basis of this interaction, we generated homology models of mTLR8 and hTLR8 bound to 2′,3′-cGMP using the agonist-bound hTLR8 structure (PDB: 3W3N)^16^ as a template (Fig. 6A). Structural comparison revealed extensive conservation of site 1 interactions. In both receptors, 2′,3′-cGMP engaged a conserved arginine (R424/R429) and aspartate (D534/D543) through hydrogen bonding, while the G ring stacked against a conserved phenylalanine (F400/F405) and was further stabilized by a conserved aspartate residue (D536/D545), forming a ternary anion-π-π interaction sandwiching the G ring. Hydrophobic contacts surrounding the G moiety were also largely conserved. In contrast, residues interacting with the ribose and phosphate groups differed between species, particularly within the LRR11-LRR12 loop that partially covers site 1 and contributes to ligand stabilization through interactions with F343/Y348. Multiple amino acid substitutions within this loop (I349- Q344, K350-Y345, G351-K346, S352-E347, and P354-L349) are predicted to alter loop conformation and ligand stabilization.

**Figure 6.**
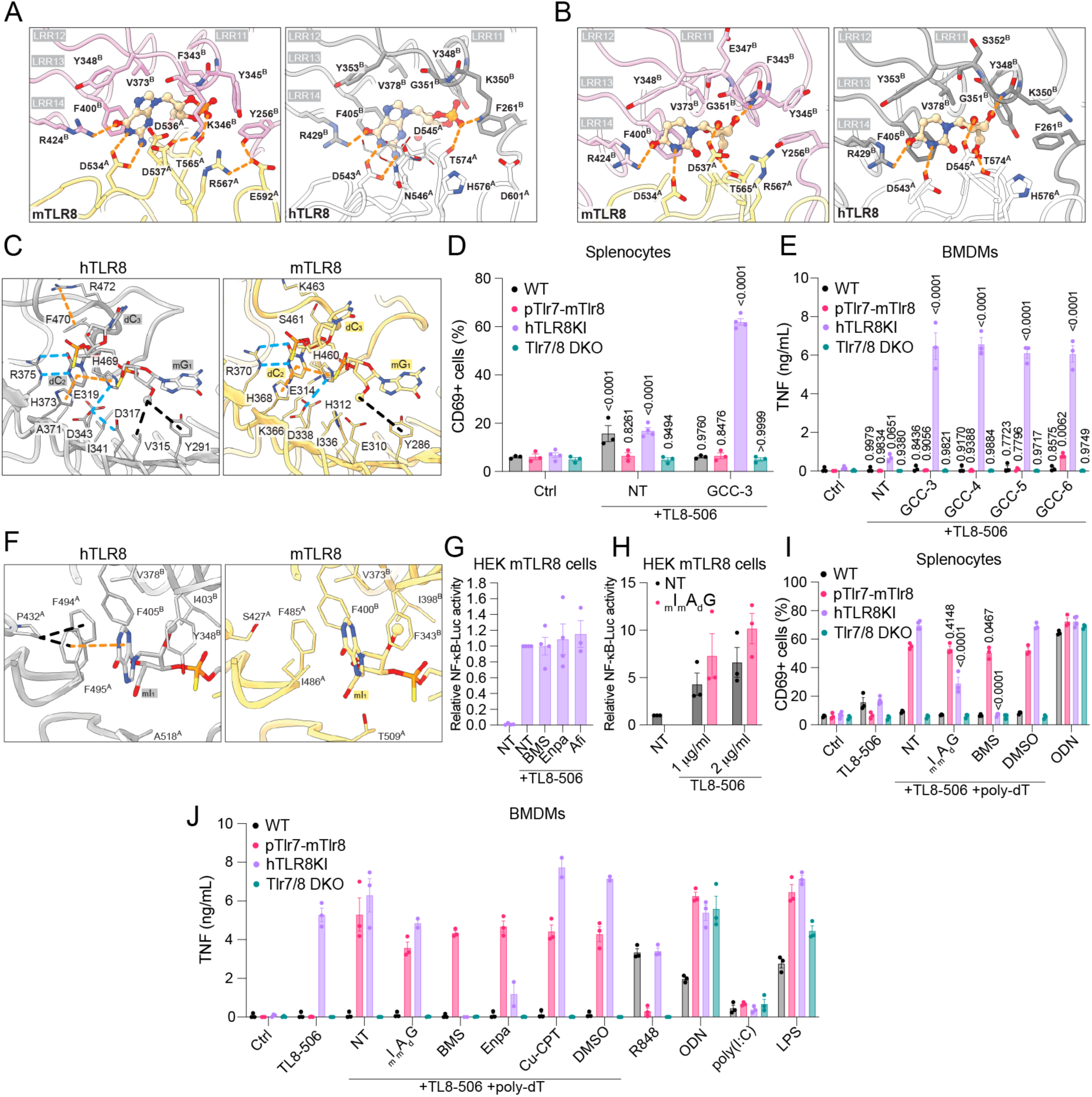
mTLR8 lacks antagonism. (**A-B**) Ligand binding site 1 in a homology model of mouse TLR8 in complex with 2′,3′-cGMP (**A,** left) or U (**B,** left). Human TLR8 structures with 2′,3′-cGMP (**A,** right) or U (**B,** right) are also shown as a comparison. Protomers of the activated mTLR8 and hTLR8 dimer are colored in pink/yellow, and white/grey respectively. The ligands 2′,3′-cGMP and U are shown in ball- and-chain representation with surrounding protein residues in stick representation. Crucial hydrogen bonds between protein and protein sidechains, or between residues are indicated in orange dashed lines. LLR11-14 are indicated. (**C**) Interactions between oligomer 38-2 and the ligand binding site 2 in human (left) and mouse (right) TLR8. Orange dashed lines indicate potential aromatic packing interactions or cation-pi interaction. Blue dashed lines represent the hydrogen bonds stabilizing the binding of oligomers at site 2. Hydrophobic interactions originating from the 2’-O-methyl group in mG_1_ are shown in black dashed lines. (**D**) Splenocytes from WT, pTlr7-mTlr8, hTLR8KI, and Tlr7/8 DKO mice were pre-treated with 5 μM GCC-3 for 30 min prior to stimulation with TL8-506 at 25 ng/mL (hTLR8KI) or 250 ng/mL (other genotypes) overnight. Single cell suspensions were stained with anti-mouse CD69 antibody and acquired on flow cytometer. Data are presented as mean of *n* = 3 mice ± s.e.m. (**E**) BMDMs from wild-type (WT), pTlr7-mTlr8, hTLR8KI, and Tlr7/8 DKO mice were pre-treated with 5 μM of GCC-3 or its length variants (GCC-4, GCC-5 and GCC-6) for 30 min prior to overnight stimulation with TL8-506 at 12.5 ng/mL (hTLR8KI) or 125 ng/mL. mTNF levels in cell supernatants were quantified by ELISA. Data are presented as mean of *n* = 3 mice ± s.e.m. (**D, E**) Two-way ANOVA with uncorrected Fisher’s LSD tests are shown compared to Ctrl conditions for each genotype. (**F**) Antagonist binding site in human TLR8 (left) and a homology model of mouse TLR8 (right) bound with GUC-v16. The loss of aromatic packing between residue 486 and guanosine base in mTLR8 site causes the lack of antagonism in mouse. (**G, H**) HEK mTLR8 cells were pre-treated with 1 μM BMS-905 (BMS), Enpatoran (Enpa), or Afimetoran (Afi), (**G**) or 5 μM _m_I_m_A_d_G^PS^ prior to overnight stimulation with 1 (**G, H**) or 2 (**H**) μg/ml TL8-506 followed by luciferase assay. Data are presented as mean of *n* = 3 independent experiments ± s.e.m. (**I**) Splenocytes from WT, pTlr7-mTlr8, hTLR8KI, and Tlr7/8 DKO mice were pre- treated for 30 min with 5 μM _m_I_m_A_d_G^PS^ or 1 μM BMS-905 (BMS) together with 1 μM poly-dT, prior to overnight stimulation with TL8-506 at 25 ng/mL (hTLR8KI) or 250 ng/mL (for other genotypes). Single cell suspensions were stained with anti-mouse CD69 antibody and acquired on flow cytometer. Data are presented as mean of *n* = 3 mice (± s.e.m. and two-way ANOVA with uncorrected Fisher’s LSD tests shown compared to “TL8-506 + poly-dT + NT” conditions for each genotype). (**J**) BMDMs from WT, pTlr7-mTlr8, hTLR8KI, and Tlr7/8 DKO mice were pre-treated for 30 min with 5 μM _m_I_m_A_d_G^PS^ or 1 μM BMS-905, Enpatoran, or Cu-CPT-8m (Cu-CPT) together with 1 μM poly-dT, prior to overnight stimulation with TL8-506 at 50 ng/mL (hTLR8KI and WT) or 125 ng/mL (pTlr7-mTlr8 and Tlr7/8 DKO), 500 nM ODN1826 (ODN), 10 μg/mL poly (I:C) or 100 ng/mL lipopolysaccharide (LPS). mTNF levels in cell supernatants were quantified by ELISA. Data are presented as mean of *n* = 3 mice ± s.e.m. _m_I_m_A_d_G^PS^, Cu-CPT8m, BMS-905, and Enpatoran conditions were conducted in *n* = 2 hTLR8KI mice.

To determine whether these differences similarly affect U recognition, we generated an mTLR8 homology model based on the hTLR8-U crystal structure (PDB: 4R0A). U adopted a binding mode in mTLR8 that was highly similar to that observed in hTLR8, forming nearly identical interactions with surrounding residues (Fig. 6B; Fig. S6A). Together, these findings support the presence of a functional site 1 agonist binding pocket in mTLR8 and suggest that its reduced responsiveness to site 1 ligands arises not from impaired ligand binding, but from structural differences in the LRR11-LRR12 loop that increase dependence on site 2 potentiators such as poly-dT (Fig. S6A).

Regardless of differences in site 1 ligand selectivity, our *in vitro* and *in vivo* data suggest that site 2 engagement synergizes with site 1 activation of mTLR8, as observed for hTLR8. However, the need for an extended _m_G_d_C_d_C ligand to elicit this effect in HEK-mTLR8 cells points to structural differences in the site 2 binding interface (see Fig. 3B). Accordingly, *in silico* modelling of mTLR8, based on the hTLR8– _m_G_d_C_d_C –U structure, identified several substitutions that weaken ligand binding. In hTLR8, Val315 and Tyr291 form a hydrophobic pocket that stabilizes the G 2′-O-methyl group, whereas replacement of Val315 with Glu310 in mTLR8 disrupts this interaction. Binding of the _m_G–_d_C phosphate is further weakened by substitution of Asp317 with His312, which also destabilizes His460 (His469 in hTLR8) and reduces its stacking interaction with the buried cytidine. In addition, replacement of Phe470 with Ser461 in the mTLR8 Z-loop likely increases loop flexibility by disrupting a cation–π interaction with Arg472 (Lys463 in mTLR8), further compromising ligand binding. Collectively, these changes are predicted to reduce the binding of mTLR8 for the _m_G_d_C_d_C trinucleotide, necessitating longer ligands for stable receptor engagement (Fig. 6C). To further compare site 2 activity between hTLR8 and mTLR8 in primary cells, we assessed site 2 responsiveness in our human and murine TLR8 mouse models using _m_G_d_C_d_C and longer derivatives in combination with TL8-506. While _m_G_d_C_d_C potentiated TL8-506 responses in hTLR8KI BMDMs and splenocytes, it had no effect in pTlr7-mTlr8, WT, or Tlr7/8 DKO cells (Fig. 6D, 6E). In contrast, a 6 nt-extended _m_G_d_C_d_C variant (i.e. _m_G_d_C_d_C_d_C_d_C_d_C) significantly enhanced TL8-506 responses in pTlr7-mTlr8 BMDMs, confirming that mTLR8 requires longer ligands than hTLR8 for effective site 2 engagement (Fig. 6E).

We recently identified a third binding site in hTLR8 that accommodates small molecules and 2′- O-methylated RNA fragments, stabilizing an open, inactive receptor conformation^19,20^. This antagonist site depends on three conserved residues present in hTLR8 (P432, F494, F495), hTLR7 (P435, F506, F507), and mTLR7 (P435, F507, F508), and mutations affecting these residues are associated with human disease^47^ (Fig. S6B). Six hTLR8 antagonist co-crystal structures (PDB: 5WYX, 5WYZ, 6KYA, 6TY5, 7YTX, and 8PFI) define a conserved pocket formed at the dimer interface and involving 25 ligand- contacting residues. Mapping sequence differences onto this pocket revealed extensive remodelling in mTLR8. Notably, substitution of hTLR8 F495 by mTLR8 I486 eliminates a key aromatic stacking interaction with 2′-O-methyl G, whereas replacement of hTLR8 P432 by mTLR8 S427 is predicted to alter the local hydrophobic architecture of the pocket (Fig. 6F; Fig. S6B). Consistent with the importance of these residues, mutations at the corresponding positions in hTLR7 (P435 and F507) markedly impaired antagonist binding *in vitro*^20^. In addition, 9 of the remaining 25 ligand-contacting residues are substituted in mTLR8 (Fig. S6C). Notably, several substitutions overlap with residues that form the ribose-facing surface of site 1 (349-352, 354 – Fig. S6A), linking divergence of antagonist and nucleoside recognition. Together, these substitutions are predicted to reshape the antagonist-binding pocket and reduce antagonist engagement in mTLR8 (Fig. 6F; Figs. S6A, S6B).

Consistent with the predicted disruption of the antagonist-binding site, several established hTLR7/8 and mTLR7 antagonists, including Enpatoran, BMS905 (Afimetoran), and the 2′-O-methyl oligonucleotide _m_I_m_A_d_G^PS^^19,48,49^ failed to inhibit TL8-506-induced activation of mTLR8 in HEK-mTLR8 cells (Figs. 6G, 6H). Similarly, _m_I_m_A_d_G^PS^ and BMS905 did not inhibit poly-dT/TL8-506-mediated mTLR8 activation in pTlr7-mTlr8 splenocytes, whereas both significantly antagonized hTLR8 signaling in hTLR8KI splenocytes (Fig. 6I). The same pattern was observed in BMDMs, where Enpatoran and BMS905 inhibited hTLR8 in hTLR8KI cells but not mTLR8-mediated responses to poly-dT and TL8-506 in pTlr7-mTlr8 cells (Fig. 6J). Notably, _m_I_m_A_d_G^PS^ and CU-CPT-8m failed to inhibit cooperative activation of hTLR8KI by poly-dT and TL8-506, despite suppressing responses to TL8-506 alone in these cells (Fig. 6J; Fig. S6D). Collectively, these findings show that Enpatoran, BMS905, and _m_I_m_A_d_G^PS^ effectively antagonize hTLR8 but not mTLR8. Together with our structural analyses identifying mutations that disrupt the antagonist-binding pocket, these results indicate that mTLR8 is largely refractory to antagonism by currently available TLR7/8 inhibitors.

## Discussion

The prevailing view that murine TLR8 (mTLR8) is non-functional arose from early observations that, unlike hTLR8 and TLR7, mTLR8 did not respond to the synthetic ligand R848 when overexpressed and that short RNA-mediated responses in mice solely depended on mTLR7^6,7,10^. These findings led to the proposal that mTLR7 functionally replaced hTLR8 in mice^6^. Subsequent reports that mTLR8 could be activated by combinations of poly-dT and site 1 agonists such as CL075 challenged this model but were difficult to reconcile with the apparent lack of activity to conventional TLR8 agonists^23,31^.

The discovery that hTLR8 utilizes multiple ligand-binding sites has fundamentally changed our understanding of TLR8 activation and provides a framework to resolve these seemingly contradictory observations on mTLR8 biology^16,18^. Here, we show that mTLR8 is not intrinsically inactive but can be directly activated by site 1 agonists, including TL8-506 and the endogenous cyclic nucleotide 2′,3′- cGMP, which we found also activates hTLR8. Consistent with previous studies, direct comparison of hTLR8 and mTLR8 ectodomains in primary cells from hTLR8KI and pTlr7-mTlr8 mice showed that mTlr8 activation by R848, uridine (U), TL8-506, and ssRNA40 was largely absent, in contrast to robust activation of hTLR8. Similarly, site 1 activity of mTLR8 was markedly weaker than that of mTLR7, particularly toward 2′,3′-cGMP, collectively suggesting that responses to isolated site 1 ligands are likely dominated by mTLR7 in mice.

As observed in our study, despite its weak response to site 1 agonists alone, compared to hTLR8 and mTLR7, mTLR8 activity was strongly enhanced by concurrent engagement of site 2. Consistent with previous observations, poly-dT potentiated TL8-506 responses in both human and mouse TLR8^23,31^. More broadly, site 2 engagement altered the selectivity landscape of mTLR8, enabling responses to ligands that were otherwise inactive. Most notably, 2′,3′-cUMP and U became specific mTLR8 agonists when combined with poly-dT, revealing a selective sensing capacity that is independent of mTLR7, and more aligned with the activity of hTLR8 (see summary of site 1, 2 activities in Fig. S7). Structural analyses further showed that the site 2 architecture of mTLR8 closely resembles that of hTLR8 while appearing more reliant on slightly longer nucleic acid ligands. Building on our recent characterization of the hTLR8 site 2 ligand _m_G_d_C_d_C, these findings support a direct role for DNA fragments in mTLR8 activation. Functionally, DNA engagement confers upon mTLR8 a unique capacity to sense bacterial DNA fragments in concert with RNA degradation products such as 2′,3′-cGMP or 2′,3′- cUMP, a mechanism independent of mTLR3, mTLR7, and mTLR9 in our assays. Comparison of ssRNA40^PS^, its U-to-A variant ssRNA41^PS^, and ORN06^PS^ revealed that ORN06^PS^ was the strongest potentiator of TLR8 activation, whereas ssRNA41^PS^ was the weakest. Given that ORN06^PS^ contains 6 GU motifs for a total of 14 U (vs 4 GU and 7 U for ssRNA40^PS^), this suggests that GU content and total U content are important determinants of RNA-mediated potentiation of TLR8 signaling. However, future studies using cell systems lacking or overexpressing DNase family members such as DNase I, DNase I-like 1/2/3, or Trex1, or RNases such as RNase T2, RNase 2, or RNase 6, will be important to directly assess the cooperative role of specific nucleases in the generation of both pathogen- and host-derived DNA and RNA ligands for modulation of mTLR8 sensing.

Crystal structures of hTLR8 ECD initially reported by Tanji et. al. showed two agonistic sites, i.e. site 1 and site 2 for U (or other small-molecule ligands) and short ssRNA fragments, respectively^18^. However, in addition to site 1 and site 2 ligands, we recently identified a third regulatory site in human TLR7, human TLR8, and mouse TLR7 that binds naturally occurring 2′-O-methyl RNA fragments and promotes an inactive receptor conformation^20^. Our data here indicate that mTLR8 differs fundamentally from hTLR8, hTLR7, and mTLR7 in its susceptibility to antagonism. Ten of the twenty-five antagonist- contacting residues are substituted in mTLR8 and are distributed across both protomers of the TLR8 dimer. Collectively, these changes extensively remodel the antagonist-binding pocket, consistent with the resistance of mTLR8 to chemically distinct inhibitors. Although additional endogenous antagonists may exist, the corresponding site is similarly disrupted in rat TLR8, which exhibits reduced responsiveness to site 1 agonists while retaining responsiveness to site 2 co-stimulation^23^. Thus, diminished site 1 sensitivity and loss of antagonist sensitivity appear to arise from a common structural mechanism. A key determinant of this divergence is in the LRR11-12 loop (residues 349-352 and 354 in hTLR8). This loop contacts the ribose moiety of site 1 nucleosides and interacts with antagonists in all currently available inhibitor-bound hTLR8 structures, yet all five residues are substituted in mTLR8. Together with Phe261, these substitutions define the ribose-facing surface of site 1. Accordingly, the same structural element that tunes site 1 ligand recognition also disrupts antagonist binding. Whether this coupling reflects adaptive evolution or relaxed selective constraint remains unclear. Murine TLR8 functions alongside TLR7 and TLR13, which may reduce evolutionary pressure to maintain high sensitivity to RNA fragments through a fully conserved site 1 architecture. However, distinguishing between these possibilities will require formal analyses of ligand selection, which have not yet been performed for rodent TLR8.

To determine whether mTLR8 contributes to physiological nucleic acid sensing, we generated mice expressing *mTlr8* under control of the endogenous *Tlr7* promoter. Replacing the promoter did not substantially alter *mTlr8* expression in BMDMs, suggesting comparable baseline expression of *Tlr7* and *Tlr8* in these cells. In contrast, splenic *mTlr8* expression was significantly increased, consistent with stronger *Tlr7* promoter activity in the spleen. As expected from previous studies demonstrating tonic TLR7 signaling in splenic B cells^45,46^, Tlr7/8 DKO splenocytes displayed reduced expression of NF-κB- associated transcriptional programs. Remarkably, increased mTLR8 expression in pTlr7-mTlr8 mice partially restored fundamental parts of this transcriptional signature, including expression of key immediate-early transcription factors such as *Junb* and *Atf3*. Expression of hTLR8 ectodomain within the same genomic context produced a similar effect and restored TLR7-dependent gene expression to near wild-type levels, consistent with the stronger activity of hTLR8. These experiments demonstrate that mTLR8, hTLR8, and mTLR7 can all respond to physiological levels of nucleic acid-derived stimuli. Nevertheless, mTLR8 failed to fully rescue the B-cell defects associated with *Tlr7-*deficiency, indicating further that its physiological activity in B cells remains substantially weaker, compared to mTLR7. Notably, the endogenous mTLR8 ligands responsible for tonic signaling in this mouse model were not identified. Using an established mouse model of TLR activation in the central nervous system, we demonstrated that intrathecal administration of the combination TL8-506 plus poly-dT induced mTLR8- dependent neuroinflammation and neurodegeneration, establishing that endogenous mTLR8 function is sufficient to drive physiologically relevant inflammatory responses independently of mTLR7.

RNase T2 and PLD3/4 have recently been shown to sequentially process endosomal RNA to release 2′,3′-cGMP, serving as optimal endogenous hTLR7 site 1 ligand, with substantially higher affinity than G^17,37,50^. Notably, exogenous 2′,3′-cGMP or G must typically be supplied at millimolar concentrations to activate TLR7 experimental systems^14,37^, and a similar trend is also observed for TLR8 activation by U^14,20^. Importantly, these experimental concentrations are unlikely to reflect physiological endosomal levels, as the endogenous concentration of 2′,3′-cGMP generated during nucleic acid degradation remains unknown. Consequently, although further studies are needed to define the impact of physiologically relevant intracellular 2′,3′-cGMP levels on innate immune responses and central nervous system injury, we view our experiments examining the effects of exogenous 2′,3′-cGMP on BMDMs and in the mouse brain as proof-of-principle studies that establish its potential biological activities.

There is rising evidence for a relevance for nucleic acid-sensing TLRs including TLR8 in various human diseases. For instance, human alleles of TLR8 with enhanced sensitivity to RNA ligands and causing severe autoimmune disease have been identified^51,52^. Aligning with this, TLR7/8 antagonists are currently under investigation as potential therapeutic concepts in autoimmune disease, e.g. Enpatoran in systemic lupus erythematosus^53^. In addition, small chemical compounds acting as agonists for nucleic acid-sensing TLRs are considered as attractive therapeutic targets in infectious disease, allergy, and cancer^54,55^. So far, mTLR8 has been largely omitted from research on nucleic acid sensing, due to the overactivity of mTLR7, presumed subordinal role of mTLR8 function, and systemic inflammation initially reported in hTLR8 transgenic mice^56^. Collectively, our data confirm that mTLR8 is functional, shares key biological features with hTLR8, and can be effectively studied in TLR7-deficient mice. However, we demonstrate that mice lacking TLR7 and expressing a humanized TLR8 ectodomain under the control of *mTlr7* promoter provides a more sensitive model, which could enable mechanistic studies of disease pathogenesis and the exploration of novel therapeutic opportunities.

To conclude, our findings establish mTLR8 as a functional nucleic acid sensor. We propose a model in which mTLR8 exhibits intrinsically weak sensitivity to site 1 ligands, which may be associated with loss of antagonist-mediated regulation through the shared LRR11-12 loop. This decreased site 1 sensitivity may limit excessive tonic activation by endogenous ligands such as 2′,3′-cGMP, otherwise potentially leading to autoimmunity as seen in mutant *Tlr7^Kika^* mice^57^. However, similar to hTLR8^14,30,34^, cooperation with site 2 ligands markedly expands mTLR8 sensing capacity, enabling robust responses to combinations of RNA and DNA fragments. Under these conditions, mTLR8 acquires ligand specificities not seen with mTLR7, including selective recognition of U and 2′,3′-cUMP combined with mixed fragments of RNA and bacterial DNA molecules. Together with the capacity of both mTLR8 and hTLR8 to partially restore tonic TLR7-dependent signaling in spleen, these findings highlight site 2 engagement as a critical determinant of TLR7/8 ligand selectivity and suggest that combinatorial nucleic acid sensing expands the repertoire of these receptors beyond site 1 ligands.

## Material and Methods Reagents

All reagents were prepared and used according to the manufacturer’s instructions, unless otherwise stated.

*Oligonucleotides (* is phosphorothioate (PS), mX is* 2’*-O-Methyl, X is DNA and rX is RNA)*

**Table 1.**
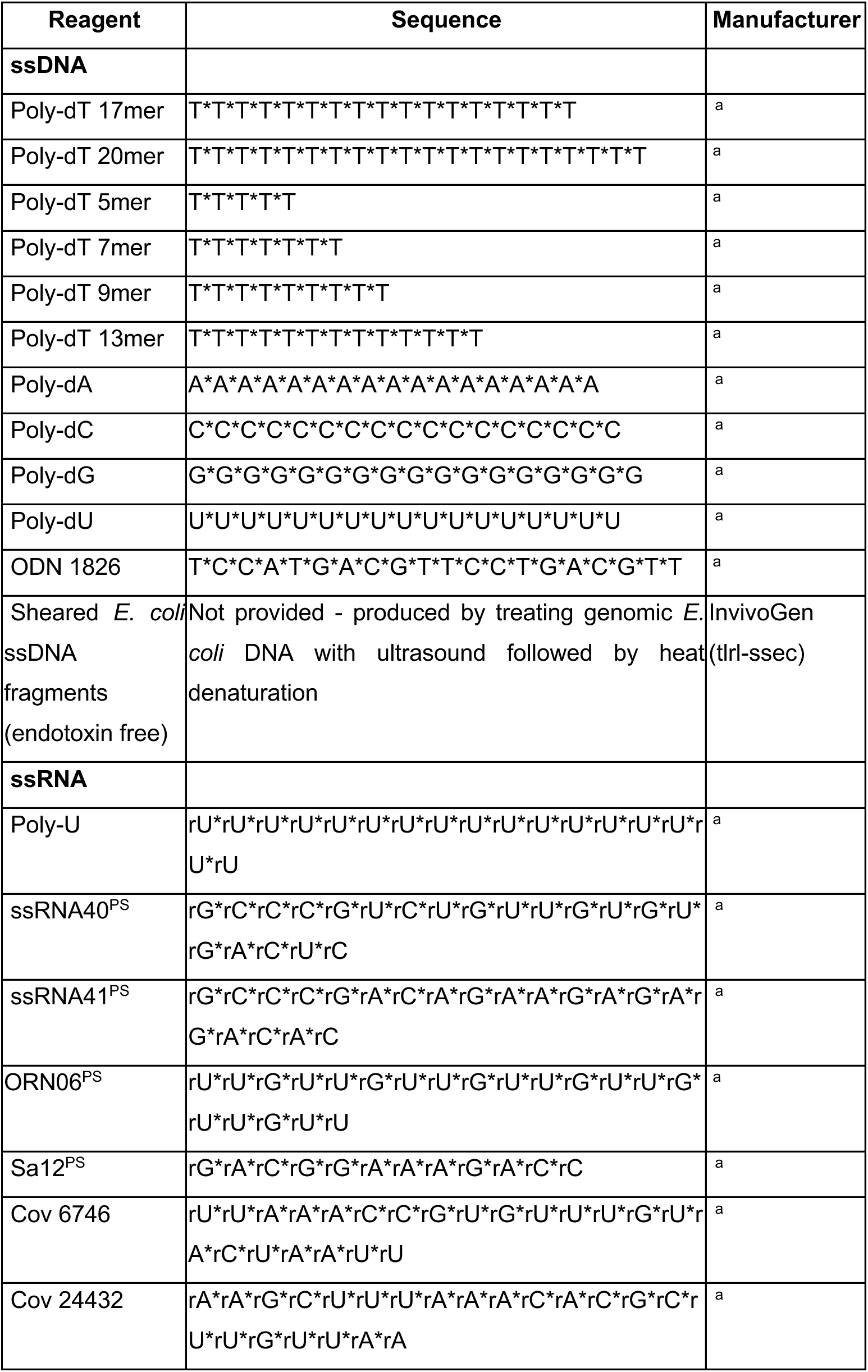
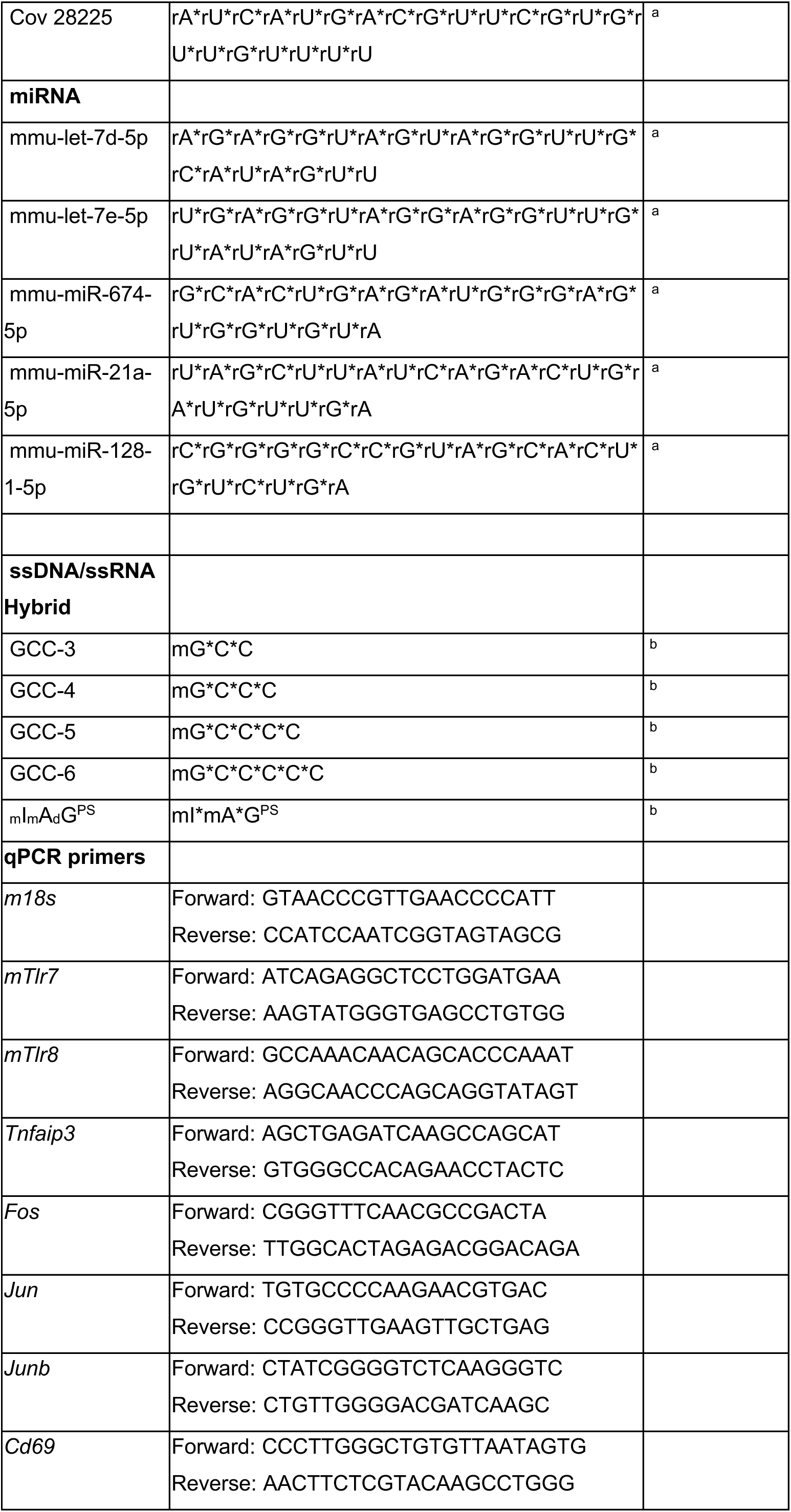
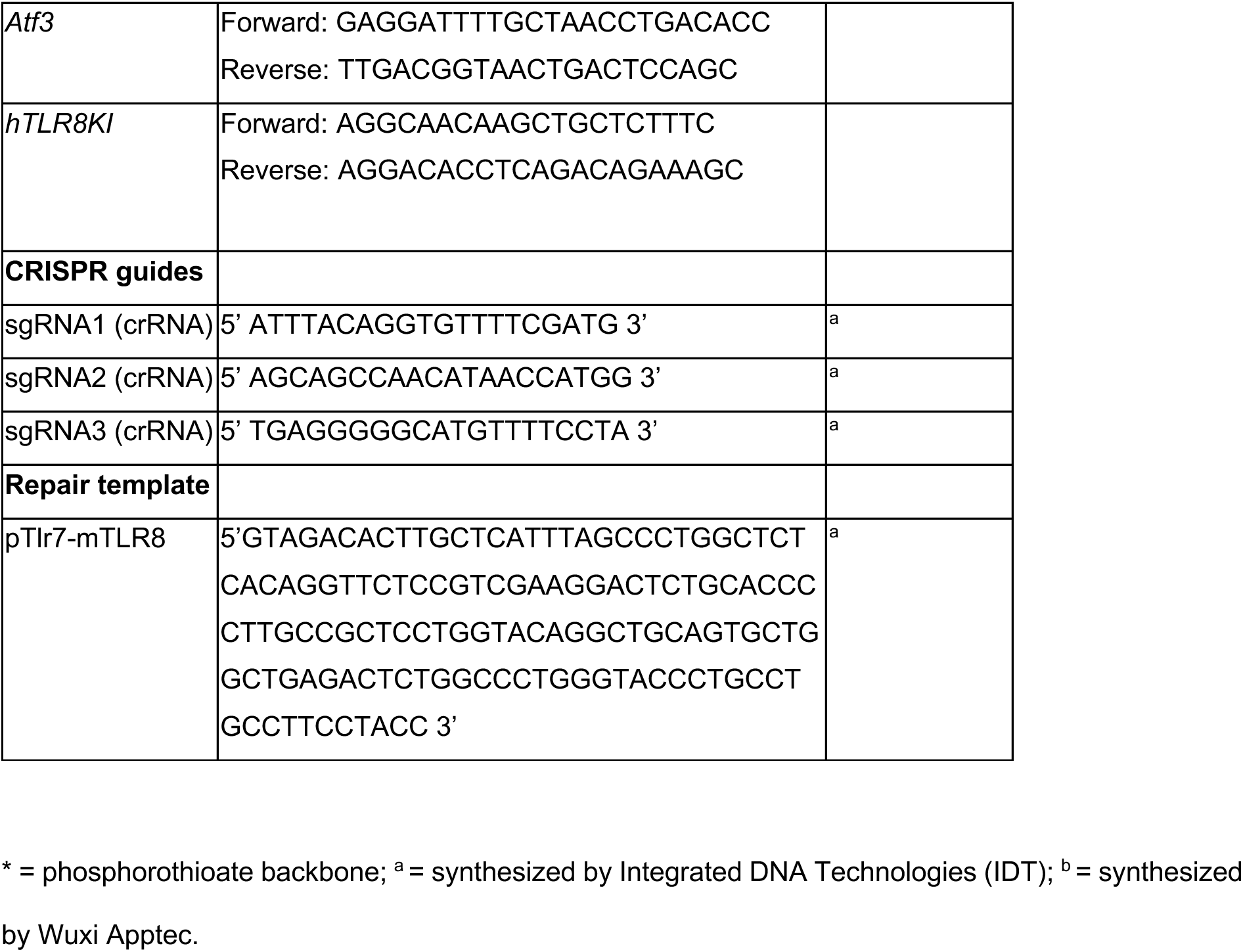
Overview of used oligonucleotides, their sequence and manufacturer.

*TLR agonists, antagonists, nucleotides, nucleosides, and others*

**Table 2.** Overview of used TLR agonists, antagonists, nucleotides, nucleosides, and manufacturer. *Preparation of reagents*

| Reagent | Catalog number | Manufacturer |
|---|---|---|
| **TLR agonists/ antagonists** |  |  |
| TL8-506 | tlrl-tl8506 | InvivoGen |
| Motolimod | HY-13773 | MedChemExpress |
| Selgantolimod | HY-109137 | MedChemExpress |
| Loxoribine | tlrl-lox | InvivoGen |
| Imiquimod | tlrl-imqs-1 | InvivoGen |
| R848 | tlrl-r848-5 | InvivoGen |
| CL075 | tlrl-c75 | InvivoGen |
| PolyIC (HMW) | tlrl-pic | InvivoGen |
| LPS | ALX-581-012 | Enzo Life Sciences |
| Enpatoran | inh-m5049 | InvivoGen |
| Afimetoran | inh-afi | InvivoGen |
| BMS-905 | HY-153254 | MedChemExpress |
| CU-CPT-8m | HY-112050-5MG | MedChemExpress |
| **Nucleosides and derivates** |  |  |
| Uridine | A15227.06 | Thermo Fisher |
| Deoxyuridine | A16026.06 | Thermo Fisher |
| Pseudouridine | SMB00912-25MG | Sigma-Aldrich |
| 5'-Methyluridine | H64259.14 | Thermo Fisher |
| 2'-O-Methyluridine | J64897.03 | Thermo Fisher |
| Uracil | U1128-25G | Sigma-Aldrich |
| Adenosine | A10781.09 | Thermo Fisher |
| Guanosine | G6264 | Sigma-Aldrich |
| Thymidine | A11493.06 | Thermo Fisher |
| Cytidine | A10261.09 | Thermo Fisher |
| Inosine | Provided by S. Bauer |  |
| Methylthioinosine<sup>36</sup> | Provided by S. Bauer |  |
| 2‘-UMP | U 015 | Biolog |
| 3‘-UMP | U 016 | Biolog |
| 5‘-UMP | U6375-1G | Sigma-Aldrich |
| 2‘-GMP | G 022 | Biolog |
| 3‘-GMP | G 021 | Biolog |
| 5‘-GMP | 226600050 | Thermo Fisher |
| 2′,3′-cGMP | G 025-50 | Biolog |
| 2′,3′-cUMP | A 307-50 | Biolog |
| 2′,3′-cAMP | U 004-50 | Biolog |
| 2′,3′-cIMP | I 032-05 | Biolog |
| 2′,3′-cCMP | C 104-50 | Biolog |
| 3',5'-cGMP | G 001-100 | Biolog |
| 3',5'-cUMP | U 001-50 | Biolog |
| **Others** |  |  |
| TNF | 300-01A | Peprotech |
| LyoVec | lyec-1 | InvivoGen |
| RNase A | EN0531 | Thermo Fisher |

### Nucleosides and nucleotides

All nucleosides and nucleotides apart from adenosine, guanosine, and xanthosine, were dissolved in nuclease-free water up to a concentration of 200 mM. Adenosine, guanosine, and xanthosine were dissolved in DMSO at a concentration of 200 mM. Subsequently, all substances were dissolved using ultrasound and then sterile-filtered through a 0.2 μM membrane. All substances except adenosine, guanosine, and xanthosine were directly added to the wells containing cells and media. Adenosine, guanosine, and xanthosine were dissolved in cell culture media in the indicated concentration. Complete dissolution of these substances was achieved by short ultrasound incubation. All dissolved nucleotides and nucleosides were sterile-filtered through a 0.2 µm filter before use.

### DNA and RNA oligonucleotides, agonists, and antagonists

All RNA oligonucleotides including ssRNAs and miRNA mimics were dissolved in nuclease-free water at a concentration of 0.8 mg/ml. Before transfection 1 volume of RNA was dissolved in 2 volumes of Lyovec transfection agent (InvivoGen, #lyec-1). DNA oligonucleotides such as ssDNAs, including poly- dT, were added to the respective cellular systems without transfection. 2’-OMe modified oligonucleotides were resuspended in Rnase-free TE buffer, pH 8.0 (Thermo Fisher Scientific) and were added directly to the cells without transfection.

## HEK Blue™ cell lines and TLR activation reporter assay

HEKBlue™ cells expressing murine/human TLR7 or murine/human TLR8 and a NF-κB/AP-1-inducible secreted embryonic alkaline phosphatase (SEAP) reporter gene, as well as the respective control cell lines HEKBlue™ Null2-k, Null1-k, Null1-v or Null1 (InvivoGen, San Diego, CA, USA) were cultured in Dulbecco’s modified Eagle’s medium (DMEM; Gibco #41965062, Thermo Fisher Scientific, Waltham, MA, USA). DMEM was supplemented with 10% heat-inactivated fetal calf serum (FCS; Gibco #10082147) and 1% penicillin/streptomycin (Gibco #15140122). In addition, selection antibiotics Zeocin (100 μg/ml; InvivoGen #ant-zn) and/or Blasticidin (10-30 μg/ml; InvivoGen #ant-bl) were used for the corresponding cell culture medium, as specified by the manufacturer. HEKBlue™ cells were seeded into 96-well plates at a density of 50,000 cells per well. SEAP reporter activity was quantified using either HEK-Blue™ Detection (InvivoGen, #hb-det2) or QUANTI-Blue™ Solution (InvivoGen), prepared according to the manufacturer’s instructions. For assays using HEK-Blue™ Detection, the cell culture medium was replaced with HEK-Blue™ Detection medium, followed by stimulation with the indicated chemical ligands, nucleosides, and/or oligonucleotides for 24 h. Alternatively, cells were stimulated in regular culture medium for 24 h, after which cell culture supernatants were transferred to QUANTI- Blue™ Solution for SEAP detection. Absorbance was measured using a SpectraMax iD3 microplate reader (Molecular Devices, San Jose, CA, USA) at 655 nm.

## HEK luciferase assay

HEK293 cells stably expressing mTLR8 (InvivoGen, San Diego, CA, USA) were reverse-transfected with pNF-κB-Luc4 reporter (Clontech), with Lipofectamine 2000 (Thermo Fisher Scientific), according to the manufacturer’s protocol. Briefly, 500,000-700,000 cells were reverse-transfected with 200-400 ng of pNF-κB-Luc4 reporter with 1.2 μl of Lipofectamine 2000 per well of a 6-well plate and incubated for 3- 24 h at 37 °C with 5% CO_2_. Following transfection, the cells were collected from the 6-wells and aliquoted into 96-wells, just before oligo and overnight TLR stimulation. The cells were then lysed in 40 μl (for a 96-well plate) of 1X Glo Lysis buffer (Promega) for 10 min at room temperature. 15 μl of the lysate was then subjected to firefly luciferase assay using 35 μl of Luciferase Assay Reagent (Promega). Luminescence was quantified with a Fluostar OPTIMA (BMG LABTECH) luminometer.

## Mice

C57BL/6 (WT), *Tlr7*^-/-^ (7KO) and *Tlr*8^-/-^ (8KO) mice were bred at the FEM, Charité – Universitätsmedizin Berlin, Germany. 7KO mice were generously provided by S. Akira (Osaka University, Osaka, Japan), while 8KO mice were generated in the Alexopoulou laboratory. TLR3/7/9 TKO (3/7/9KO) mice were generated and bred at the Bauer laboratory, as previously described^58^. C57BL/6, 7KO and 8KO animals were maintained at the Charité – Universitätsmedizin Berlin according to the guidelines of the committee for animal care, and animal procedures were approved by the Landesamt für Gesundheit und Soziales (LAGeSo) Berlin, Germany.

*Tlr7/8* double-deficient (Tlr7/8 DKO), hTLR8KI, and pTlr7-mTlr8 mice were produced via CRISPR genome editing by the Monash Genome Modification Platform, Monash University (Clayton, Australia) as a node of Phenomics Australia. Breeding of these colonies and splenocyte experiments were approved by Monash Medical Centre B Animal Ethics Committee (#MMCB/2024/18/BC and #MMCB/2024/30). Tlr7/8 DKO, hTLR8KI, and pTlr7-mTlr8 mice were housed in solid bottom cages (Tecniplast 1264C IVC) containing autoclaved pine sawdust bedding and provided with environmental enrichment. Animals were maintained under specific pathogen-free conditions in a temperature- controlled (18-24°C) and humidity-controlled (40-70% relative humidity) facility on a 12 h light/dark cycle with 25-50 air changes per hour. Mice had ad libitum access to irradiated rodent chow (Specialty Feeds) and autoclaved tap water.

## Tlr7/8 DKO and hTLR8KI mouse generation

Cas9 nuclease (50 ng/ml - IDT Alt-R® S.p. HiFi Cas9 Nuclease V3, IDTDNA) was incubated with the sgRNAs comprised of both crRNA and tracrRNA sequences (sgRNA1 and sgRNA2) (IDTDNA) to form a ribonucleoprotein (RNP) complex. A 2878 bp human TLR8 ectodomain ssDNA repair template (Fig. S8) was generated using the Guide-it Long ssDNA Production System (Takara) according to the manufacturer’s instructions. Cas9 nuclease (80 ng/μl) was incubated with the sgRNAs (80 ng/μl) to form a ribonucleoprotein (RNP) complex. The RNP and the ssDNA repair template (60 ng/μl) were microinjected into C57BL/6J zygotes at the pronuclei stage. Microinjected zygotes were transferred into the uterus of pseudo pregnant F1 females. Positive mice were back-crossed twice on C57BL/6J background to obtain N2 generation progeny used in these experiments. Notably we obtained two different Tlr7/8 DKO lines lacking insertion of the repair template (therefore lacking Tlr7 and most of the Tlr8 locus); one line contained a 508 bp random insert. Both lines were found to be non-functional for mTlr7 and mTlr8 and were phenotypically identical. Except for the RNA sequencing data and RTqPCR validations in Fig. 5I, males from the Tlr7/8 DKO line with the 508 bp random insert were used.

## pTlr7-mTlr8 mouse generation

Unlike the other lines above, this line was generated to preserve expression of the mTlr8 gene, placed under the control of mTlr7 promoter. The same sgRNA1 was used to cut after the mTlr7 initiating ATG, however a different sgRNA3 was used to cut upstream of mTlr8. Cas9 nuclease (IDT Alt-R® S.p. HiFi Cas9 Nuclease V3, IDTDNA) was incubated with the sgRNAs comprised of both crRNA and tracrRNA sequences (IDTDNA) to form an RNP complex. Cas9 nuclease (3.8 μM), the sgRNA (4.2 μM) and the repair template [pTlr7-mTlr8] (10 μM) were electroporated into C57BL/6J zygotes at the pronuclei stage. Electroporated zygotes were transferred into the uterus of pseudo pregnant F1 females. Positive mice were back-crossed thrice on C57BL/6J background to obtain N3 generation progeny used in these experiments.

## Intrathecal injection into mice

Intrathecal injections into mice were performed as described previously^41^. Briefly, male 6-8-week-old C57BL/6, 7KO, and 8KO mice from various litters received intrathecal injections of 276 µg 2′,3′-cGMP, corresponding to a theoretical final intrathecal concentration of 10 mM, or 1 µg TL8-506 in combination with 0.8 nmol phosphorothioate-backboned 17-mer poly(dT), corresponding to theoretical final intrathecal concentrations of 12.5 µg/mL TL8-506 and 20 µM 17-mer poly(dT), respectively. Nuclease- free water was used as vehicle control. The total injection volume was 40 µL for all conditions. Final intrathecal concentrations were calculated assuming an approximate cerebrospinal fluid (CSF) volume of 40 µL and complete mixing of the injected solution with the CSF. Subsequently, mice were sacrificed 72 h after injection. After transcardial perfusion with 4% paraformaldehyde (PFA), brains were removed and subsequently cryoprotected with sucrose (10-30%) over a course of 3 d. Representative brain sections (level 1: interaural 6.60 mm; level 2: 5.34 mm; level 3: 3.94 mm; level 4: 1.86 mm; level 5: −0.08 mm) were placed on 12-mm-glass cover slips in a 24-well-plate, fixed with 4% PFA (200 μl), washed 3x with 200 μl PBS, and treated with blocking solution (5% normal goat serum (NGS); 0.2% TritonX-100) for 1 h. Immunostaining was performed, as described previously^41^. For primary antibodies, we used antibodies to NeuN (1:1,000, clone A60, Millipore) and Iba1 (1:1,000, cat. #019-19741, Wako). DAPI was obtained from Roche. Fluorescence microscopy was performed on an Olympus BX51 microscope and on a confocal laser scan Leica TCS SL microscope with sequential analysis (argon, 488 nm; helium neon, 543 nm). Stereological quantification of microglial cells and neuronal loss in mice was performed using the CellSense software provided by Olympus. Four fields (×60) in the cortex of the right hemisphere of five representative brain sections stained with Iba1 or NeuN antibody per animal were analyzed by quantifying Iba1^+^ and NeuN^+^ objects, as indicated. The threshold was set to 400 dpi.

## Primary culture of murine BMDMs

Bone marrow-derived macrophages (BMDMs) were generated from 10-16-week-old male C57BL/6J wild-type (WT), 7KO, 8KO, and 3/7/9KO mice, and BMDMs from 6-12-week-old male Tlr7/8 DKO, pTlr7- mtlr8 promoter, and hTLR8KI mice. Bone marrow was flushed from the femurs using complete DMEM and cultured in DMEM supplemented with 10% fetal bovine serum, antibiotics, and 20% L929 cell- conditioned medium as a source of M-CSF. Cells were incubated at 37 °C in 5% CO₂. On day 3, the culture medium was replenished with fresh complete DMEM and L929 cell-conditioned medium, and non-adherent cells were recovered and returned to the cultures. Cells were further differentiated until day 6, when adherent macrophages were harvested by scraping and subsequently plated for experimental stimulation.

## Tumor Necrosis Factor Enzyme-Linked Immunosorbent Assay

Primary murine BMDMs were seeded in 96-well plates at 80,000 cells per well and stimulated overnight with the indicated chemical compounds at the indicated concentrations. Cell culture supernatants were subsequently collected and stored at −70 °C until analysis. For 7KO, 3/7/9KO and 8KO BMDM figures, TNF concentrations in the supernatants were quantified using a mouse TNF enzyme-linked immunosorbent assay (TNF ELISA) according to the manufacturer’s instructions (TNF alpha Mouse Uncoated ELISA Kit, Invitrogen, #88-7324-88, Carlsbad, CA, USA). For Figures with Tlr7/8 DKO, pTlr7- mTlr8, and hTLR8KI BMDMs, mouse TNF levels were measured using TNF (BD Biosciences, #550534), and Tetramethylbenzidine substrate (Thermo Fisher Scientific) was used for quantification of the cytokines on a Fluostar OPTIMA (BMG LABTECH) plate-reader.

## Multiplex immunoassay

Primary murine WT, 7KO, or 8KO BMDMs were seeded in 96-well plates at 80,000 cells per well and stimulated for 16 h with LPS (100 ng/mL), poly(I:C) (10 μg/mL), R848 (1 μg/mL), TL8-506 (500 ng/mL), ODN1826 (500 nM), TL8-506 in combination with poly-dT (5 µM), or 2′,3′-cGMP (10 mM) alone or in combination with poly-dT. Untreated cells served as controls. Following stimulation, cell culture supernatants were collected, and cytokine (GM-CSF, IL-6, IL-10, IL-12p70, KC/GRO, VEGF-A, IL-23, IL-27p28/IL-30, IFN-β, MMP-9, IP-10, MCP-1, MIP-2, MIP-1β, NGAL/LCN2) concentrations were determined using a custom U-PLEX multiplex immunoassay (Meso Scale Diagnostics, Rockville, MD, USA) according to the manufacturer’s instructions. Briefly, capture antibodies were coupled to the corresponding U-PLEX linkers and immobilized on U-PLEX plates. Three plates with different sample dilutions were used: 4-fold: GM-CSF, IL-6, IL-10, IL-12p70, KC/GRO, VEGF-A, IL-23, IL-27p28/IL-30, IFN-β, MMP-9. 20-fold: IP-10, MCP-1, MIP-2, MIP-1β. 100-fold: NGAL/LCN2. Standards were added to the plates, followed by incubation with the respective SULFO-TAG–conjugated detection antibodies. After washing, MSD Read Buffer was added, and electrochemiluminescence signals were measured using a MESO SECTOR S 600 instrument (Meso Scale Diagnostics). Cytokine concentrations were calculated from the respective standard curves.

## Splenocyte purification, stimulation, and flow cytometry analysis

Splenocytes were purified from fresh spleens by passing through a 70 μM filter and treating with Red Blood Cell Lysing Buffer Hybri-Max (Merck) for 5 min at RT. Lysis reactions were neutralized with complete DMEM and cells were pelleted (300 × *g* for 5 min) prior to lysis for RNA purification or resuspension in medium for plating. Purified splenocytes were plated at ∼1.5x10^6^ cells per well in 24- well plates and stimulated overnight with respective agonists (25-500 ng/mL TL8-506; 1 μg/mL of R848; 20 mM Uridine; 500 nM of TLR9 agonist ODN1826; 100 ng/mL of TLR4 agonist LPS). For mTLR8 activation, TL8-506 was used after 30 min pre-treatment with 1 μM 20-mer poly-dT. BMS905 and _m_I_m_A_d_G^PS^ were co-incubated with 20-mer poly-dT for 30 min prior to overnight stimulation with TL8-506 and 20 mer-poly-dT. Cells were subsequently pelleted and rinsed with FACS buffer (2% FCS and 1 mM EDTA in 1x DPBS), then incubated for 15 min at 4°C with Purified Rat Anti-Mouse CD16/CD32 (Mouse BD Fc Block™) Clone 2.4G2 (RUO) (#553141 BD Biosciences), diluted 1:500 in ∼100 μL final volume. Cells were then stained for 30 min with ABflo® 647 Rabbit anti-Mouse CD69 mAb (#A24939-200, Abclonal); unstained cells were included along with cells stained with the matching isotype control, ABflo® 647 Rabbit IgG isotype control (#A22070-100, Abclonal). Cells were rinsed with FACS buffer and stained with DAPI (#MBD0015-1ML Sigma-Aldrich), diluted 1:500, prior to acquisition on a BD LSR- Fortessa X-20 Flow Cytometer using the BD FACS Diva software, with data analysis using FlowJo v10.9. Gating strategy is provided in Fig. S9.

## RNA and RT-qPCR analyses

Total RNA was purified from primary BMDMs or primary splenocytes using the PureLink RNA Mini Kit (Thermo Fisher Scientific) and DNase-treated using the Purelink DNASE set (Thermo Fisher Scientific). Random hexamer cDNA was synthesised from isolated RNA using the High-Capacity cDNA Archive kits (Thermo Fisher Scientific) according to the manufacturer’s instructions. RT-qPCR was carried out with the Power SYBR Green Master Mix (Thermo Fisher Scientific) on a QuantStudio 6 Flex RT-PCR system (Thermo Fisher Scientific) with the QuantStudio™ Real-Time PCR Software v1.7.2. Each PCR was performed in technical duplicate and mouse 18S was used as the reference gene. Each amplicon was gel-purified and used to generate a standard curve for the quantification of gene expression. Melting curves were used in each run to confirm specificity of amplification. Primer sequences are provided in the oligonucleotides table above.

## Flow cytometer analyses of splenic B cell subsets

For the analyses shown in Fig. S5E, splenocytes were isolated from fresh spleens using passing through 70 μM filter and treated with Red Blood Cell Lysing Buffer (155 mM NH_4_Cl, 10 mM KHCO_3_ and 0.1mM Na_2_EDTA, pH7.3) for 3 min. Cells were pelleted by centrifugation and washed twice with FACS buffer prior to staining prior to plating. 5 x 10^6^ splenocytes were incubated with TruStain FcX™ anti-mouse CD16/32/64 antibody (BioLegend, #422302) and eBioscience™ Fixable Viability Dye eFluor™ 780 (Invitrogen, #65-0865-14) diluted in PBS for 20 min at room temperature in the dark, followed by incubation with antibody cocktails diluted in Brilliant Stain Buffer containing: for PB and ABCs anti-mouse B220 BUV737 (#103248, BD Biosciences) anti-mouse CD19-BUV395 (#563557, BD Biosciences), anti- mouse CD21/35-PE, anti-mouse CD11b Percp Cy5.5 (#), anti-mouse CD23-BV421, anti-mouse CD3- BV510 (#100234, Biolegend), anti-mouse IgM-FITC (#553437, BD Biosciences), anti-mouse CD11c- A647 (#117312, Biolegend), anti-mouse CD138-BV605 (#142515, Biolegend), anti-mouse CD98-PE- Cy7 (#128214, BD Biosciences); for GC B cells anti-mouse B220 BUV737 (#103248, BD Biosciences) anti-mouse CD19-BV510 (#115546, Biolegend), anti-mouse CD95-PE (#554258, BD Biosciences). Cells were fixed and permeabilised using the eBioscience™ Foxp3/Transcription Factor Staining Buffer Set (Invitrogen, #00-5523-00), and the GC B cell panel incubated with anti-mouse/human Bcl6-A647 (#561525, BD Biosciences) diluted in 1X Perm Buffer. Samples were acquired on a BD LSRFortessa™ or BD LSRFortessa™ X-20 flow cytometer using the BD FACS Diva software, with data analysis using FlowJo v10.9. These analyses were performed blinded.

## Bulk RNA-sequencing of primary mouse splenocytes

Libraries were generated using an in-house multiplex RNA-seq method (Version 01/03/2024; Hudson Genomics Facility). 25 ng of total RNA from each sample was tagged with a 10 bp sample index and 10 bp unique molecular identifier (UMI) during initial poly(A) priming and pooled samples amplified using template switching oligo. After cDNA amplification Illumina P5 adaptors with distinct i5 indexes were added by tagmentation by Nextera transposase and PCR. Final libraries were pooled for subsequent processing and single-end sequencing was performed on a NextSeq 2000 run using a P2 100cycle kit. For sequencing, the multiplex library was loaded at 750pM with ∼1% PhiX for on-board denaturation and clustering. Base calling was performed using Dragen BCLConvert (v4.2.7). Demultiplexing of raw sequencing files was performed by the sequencing facility. A minimum of 14.9 million reads were assigned to each demultiplexed sample.

## RNA-seq analysis

RNAseq read alignment and UMI counting was performed in R (https://www.R-project.org/; v4.3.3). The scPipe package (v1.14.0)^59^ was employed to process the data. Read alignment was performed on R1 FASTQ files using the Rsubread package (v2.6.4)^60^. An index was built using the Ensembl *Mus musculus* GRCm39 primary assembly genome file and alignment was performed with default settings. Aligned reads were mapped to exons using the sc_exon_mapping function with the Ensembl *Mus musculus* GRCm39 GFF3 genome annotation file (v115).

Reads mapping to exons in the resulting BAM files were associated with each individual sample using the sc_demultiplex function, taking the UMI into account, and an overall count for each gene for each sample was generated using the sc_gene_counting function (with UMI_cor = 1). A minimum of 88% of reads per sample mapped to the reference genome, with between 50–64% of reads mapping to exons. A proportion of sequencing reads for one sample (WT_119_Spleen) had a single base mismatch in the barcode sequence. To account for this, the mismatched barcode was manually removed from the ’index.csv’ file generated by the sc_detect_bc function before running sc_demultiplex. Subsequent analyses were performed in R (v4.5.0).

Additional gene annotation was obtained using the biomaRt package^61^ (v2.66.0), a DGEList object was created with the counts and gene annotation using the edgeR package^62^ (v4.8.0), and a design matrix was constructed, incorporating mouse genotype. Lowly expressed genes were removed using the filterByExpr function and normalisation factors were calculated using the TMM method^63^. Library sizes after filtering ranged from ∼1.8–2.3 million gene counts, with between ∼16,300–16,700 individual genes detected. Counts were transformed using the voom method^64^ and a linear model was fit using the edgeR voomLmFit function, with mouse ID included as a blocking factor to account for differences based on individual variability. Differential gene expression analyses were performed using the limma (v3.66.0) package^65^. Groups were compared using the contrasts.fit function and empirical Bayes moderated *t-*statistics were calculated using the eBayes function^66^. Differentially expressed genes were determined using a false discovery rate (FDR) adjusted *p*-value < 0.05.

Heat maps were made using the pheatmap package (v1.0.13; https://CRAN.R-project.org/package=pheatmap). Hallmark gene sets were obtained from the Broad Institute Molecular Signature Database^67,68^, via the msigdbr package (v25.1.1). Log_2_ counts per million (CPM) expression values were calculated using the edgeR cpm function. Relative log_2_ CPM expression values were calculated for each gene by subtracting the average log_2_ CPM expression value in wild type controls.

Where relevant, log_2_ fold changes between different groups were also shown, indicating adjusted *p*- values as \**p* < 0.05, \*\**p* < 0.01 and \*\*\**p* < 0.001. Venn diagrams were created using the vennDiagram function from limma. Heat map scales were truncated where indicated in figure legends. The original demultiplexed R1 FASTQ files were deposited along with UMI-adjusted counts in the NCBI Gene Expression Omnibus as a SuperSeries with accession GSE349267.

## Structural contact analysis and cross-species mapping

*Structures.* All publicly deposited X-ray structures of the human TLR8 ectodomain with a bound site 1, site 2 or antagonist-site ligand were analyzed (*n* = 13), as deposited and without re-refinement. Agonist and site 2 complexes: 3W3J (CL097), 3W3K (CL075), 3W3L and 3W3N (R848), 4R07 and 4R08 (uridine and a UG dinucleotide, from ORN06 and ssRNA40 respectively) and 4R09 (uridine, a UG dinucleotide from ORN06S, and a uridine 2′,3′-cyclic monophosphorothioate). Antagonist complexes: 5WYX (CU- CPT8m), 5WYZ (CU-CPT9b), 6KYA (TH1027), 6TY5 (Compound 11), 7YTX (E6742) and 8PFI (MHV370). Resolutions range from 2.0 to 2.9 Å.

*Contacts.* A residue was scored as contacting a ligand when any of its heavy atoms lay within 5.0 Å of any heavy atom of that ligand; hydrogens were ignored throughout. Every qualifying atom pair was recorded with its distance and with the protein atom classified as main-chain or side-chain. Where an entry contains more than one crystallographically independent copy of the receptor, contacts were computed on a single dimer, so that every reported contact is an observation in one receptor dimer rather than an average. Contacts were then pooled across entries: support for each residue is given as k/N, the number of entries (k) in which the contact is observed out of those containing a ligand at that site (N), together with the closest approach seen. All distances are quoted to two decimal places at most, and differences below 1 Å between independent copies were not interpreted.

Ligand decomposition. To separate residues that read the nucleobase from those that read the ribose, nucleoside and nucleotide ligands were partitioned into phosphate, sugar and base groups by traversing the covalent bond graph of the deposited chemical component, rather than by atom-name convention; this makes the partition valid for monolithic non-standard components, where atom naming is unreliable. Each contact inherits the group of its ligand atom, and a residue may therefore contact more than one group.

*Inter-residue geometry.* Distances quoted between two protein residues (Pro432 to Phe494 and Phe495; Asp545 to His576; Asp317 to Glu319) were measured directly on the deposited coordinates across every protomer present in the asymmetric unit, using the closest heavy-atom approach between the relevant side chains, and are reported as the observed range.

Cross-species mapping. Human, mouse and rat residue equivalences were derived by pairwise global (Needleman–Wunsch) alignment of the full-length sequences with Biopython 1.87 (PairwiseAligner), using BLOSUM62 and gap penalties of −11 for opening and −1 for extension, the NCBI BLASTP defaults. Sequences were UniProt Q9NR97 (human TLR8, reviewed), P58682 (mouse, reviewed) and A0ABK0LFG4 (rat, unreviewed). A position was called substituted when the aligned residue identity differs from human, or where a human residue aligns to a gap. Offset arithmetic was not used at any point, because the offset is not constant: an indel in the Z-loop makes the human-to-mouse offset −5 up to approximately residue 432 and −9 from approximately residue 469, and human Arg438– Ala442 has no rodent counterpart.

*Limits of the rodent comparison.* No experimental structure of mouse or rat TLR8 has been determined. Every statement about a rodent residue is therefore a sequence-derived assignment mapped onto human coordinates and is reported as such.

## Language editing

Microsoft Copilot was used to assist with language refinement and improve readability. All scientific content, interpretation, and conclusions were generated and verified by the authors.

## Statistics

Data are expressed as mean ± s.e.m. Statistical analyses were carried out using Prism 10 (GraphPad Software Inc.). The exact sample size (*n*) for each experimental group/condition is provided in each figure legend and measurements shown were from distinct samples (in bar graphs, each dot represents data obtained from an independent experiment or an individual mouse). One-way and two-way analyses of variance (ANOVA) with uncorrected Fisher’s LSD were used when comparing groups or pair of conditions.

## Resource availability Lead contact

Requests for further information and resources should be directed to and will be fulfilled by the lead contacts, Seija Lehnardt and Michael Gantier.

## Materials availability

Mice or sperm from pTlr7-mTlr8 and Tlr7/8 DKO mice generated in this work is available upon direct request to M.P.G.

## Data availability

The main data supporting the results in this study are available within the paper and its Supplementary Information. Source data for the figures will be provided with this paper. RNA sequencing data has been deposited in the NCBI Gene Expression Omnibus (GEO) with accession GSE349267. Further inquiries can be directed to the corresponding authors.

## Acknowledgments

We thank J. Rientjes (Monash Genome Modification Platform, Australia) for designs of pTlr7-mTlr8 and hTLR8KI mice; J. Treloar, A. Price, L. Saunders, E. Burrowes and E. Vargas from the MHTP Animal Facility (Hudson Institute, Australia) for maintenance of the pTlr7-mTlr8, Tlr7/8 DKO, and hTLR8KI mice; O. Laczka for feedback on the manuscript. We thank the Lehnardt lab for helpful discussions. This work was supported by Deutsche Forschungsgemeinschaft (DFG) LE 2420/2-1, SFB-TRR167 (“NeuroMac”)/B03, SFB-TRR1315 (“Mechanisms and Disturbances in Memory Consolidation: from synapses to systems”)/A01 (to S.L.); the Australian National Health and Medical Research Council Project Grant (grant number 2020565 to M.P.G., J.I.E and B.C.); mRNA Victoria Research Acceleration Fund, the Victorian Government’s Operational Infrastructure Support Program and COVID-19 Treatments Medical Research Fund (M.P.G.); Scholarships by the Einstein Center for Neurosciences Berlin (to V.K. and H.M.),and the Sonnenfeld-Stiftung (to M.B.). G.K. and P.S. were supported by the DFG through grants of the SFB1423 “Structural Dynamics of GPCR Activation and Signaling”, project number 421152132, subprojects A01/Z03 (to P.S.); through SFB1078 “Protonation Dynamics in Protein Function”, project number 221545957, subproject B06 (to P.S.), and Germany’s Excellence Strategy—EXC 311 2008/1 (UniSysCat)—390540038 (Research Unit E) (to G.K., P.S.). E.L. is funded by the European Union’s Horizon 2020 research and innovation programme under grant agreement No. 848146 (To_Aition), the state of Berlin and the European Regional Development Fund through the grant EFRE PersMedLab, the Deutsche Forschungsgemeinschaft (DFG, German Research Foundation) under Germany’s Excellence Strategy – EXC2151 – 390873048 “ImmunoSensation3 - the immunosensory system: from immunodiversity to precision medicine”, DFG SFB1454 – 432325352 “Metaflammation and Cellular Programming”, SFB1403 – 414786233 “Cell Death in Immunity, Inflammation and Disease”, TRR237 – 369799452 “Nucleic Acid Immunity” and GRK2168 – 272482170 “Myeloid antigen-presenting cells and induction of adaptive immunity”. E.R. was supported by a Research Training Scholarship from Monash University. We acknowledge use of equipment and technical support from RNAte (Hudson Institute of Medical Research, Clayton, Australia). Generation of the novel mice described here was conducted by a node of Phenomics Australia, which is supported by the Australian Government Department of Education through the National Collaborative Research Infrastructure Strategy, the Super Science Initiative and the Collaborative Research Infrastructure Scheme. This research project was undertaken with the assistance of resources and services from the National Computational Infrastructure and the Pawsey Supercomputing Research Centre, which are supported by the Australian Government and the Government of Western Australia accessed through the National Merit Allocation and ANU Merit Allocation schemes.

## Author contributions

Conceptualization, S.L., M.P.G., L.H., E.R., M.B.; Formal analysis, L.H., E.R., M.B., Q.S., A.L.M., R.J., S.H., J.B., B.C., C.C., J.I.E., G.K., V.K., P.B., and H.M.; Funding acquisition, S.L., M.P.G., J.I.E., E.L., G.K., B.C., P.S., and G.K.; Investigation, L.H., M.B., E.R., W.S.N.J., S.S., Q.S., A.L.M., R.J., S.H., J.B., C.C., D.S.S., A.K., V.K., H.M., C.H., and C.K.; Supervision, S.L., S.B., E.L., and M.P.G.; Visualization, L.H., M.B., E.R., M.P.G., G.K., and P.S.; Material, D.G., L.A., and S.B.; Writing - original draft, L.H., M.B., S.L., M.P.G., and S.B. with support of all co-authors. All authors have read and agreed to the published version of the manuscript.

## Supplementary Information

Supplementary Figures S1-9.

---

**Supplementary Figure 1.**
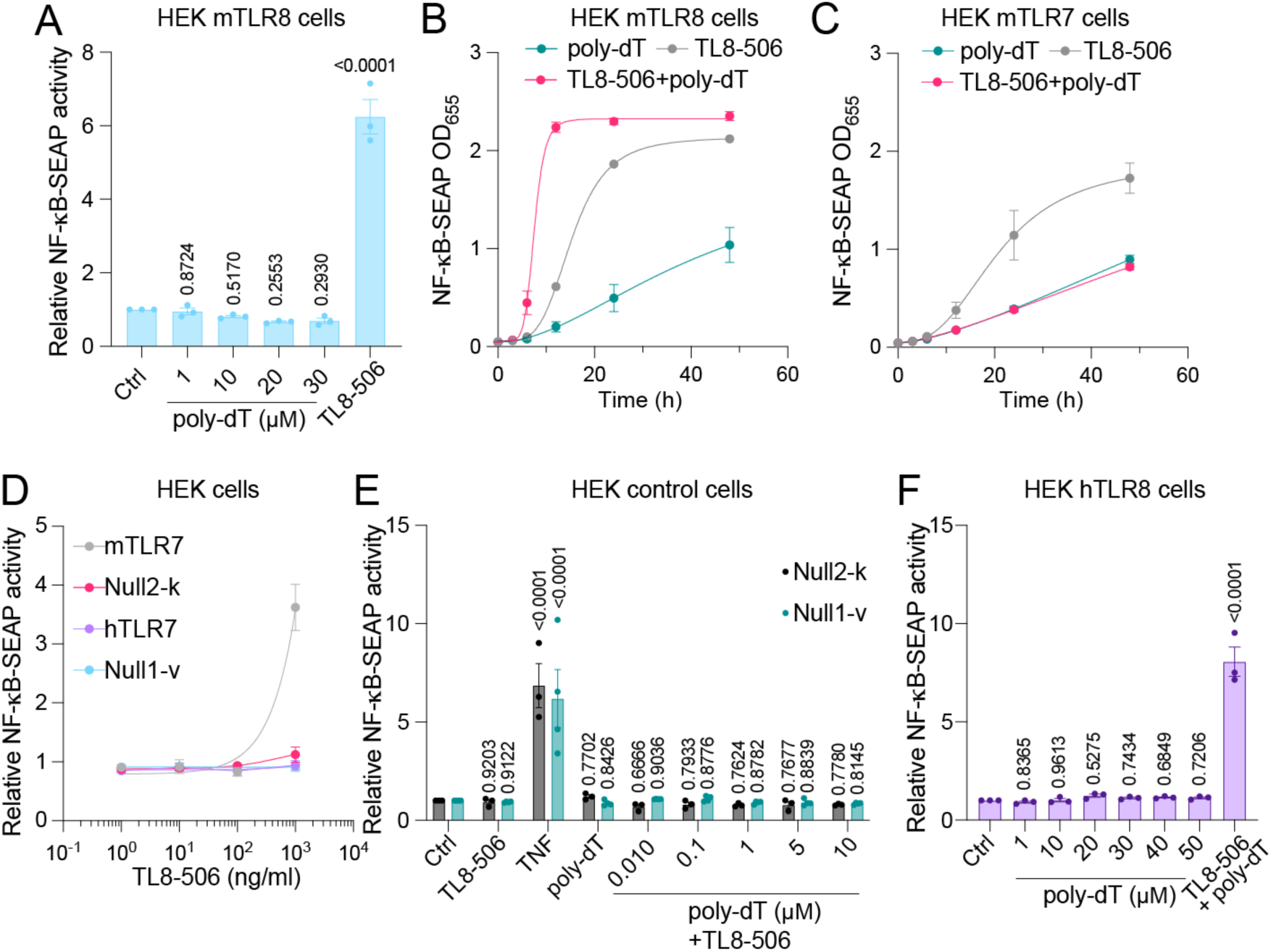
Response of HEK mTLR8, mTLR7, and hTLR8 reporter cells and parental control cell lines to TL8-506. (**A**) HEK-Blue cells co-expressing murine TLR8 (mTLR8) and an NF-κB/AP1-inducible secreted embryonic alkaline phosphatase (SEAP) reporter gene were incubated with TL8-506 (500 ng/ml) or with increasing deoxythymidine 17-mer (poly-dT) doses, as indicated, for 24 h. HEK mTLR8 (**B**) or mTLR7 (**C**) reporter cells were incubated with TL8-506 (500 ng/ml), alone or in combination with poly-dT (5 µM), for various time periods, as indicated. (**D**) HEK mTLR7, HEK hTLR7, (parental line-derived) Null2-k, or Null1-v cells were incubated with increasing TL8-506 doses, as indicated, for 24 h. (**E**) Null2-k or Null1-v were incubated with TL8-506 (500 ng/ml) or 17-mer poly-dT (5 µM), alone or in combination, as indicated. TNF (100 ng/ml) was used as positive control for SEAP induction. (**F**) HEK human TLR8 (hTLR8) reporter cells were incubated with various concentrations of 17-mer poly-dT, as indicated, or with the combination of TL8-506 (500 ng/ml) and poly-dT (5 µM), for 24 h. (**A-E**) Subsequently, optical density was assessed. Unstimulated cells (Ctrl) served as negative control. Data are mean of *n* = 3 (A, E [Null2-k], F) or *n* = 4 (B, C, D, E [Null1-v]) independent experiments. Data are shown as fold change of the optical density normalized to unstimulated cells (Ctrl) (± s.e.m. and [A, F] one-way or [E] two-way ANOVA with uncorrected Fisher’s LSD tests shown comparing to Ctrl conditions).

**Supplementary Figure 2.**
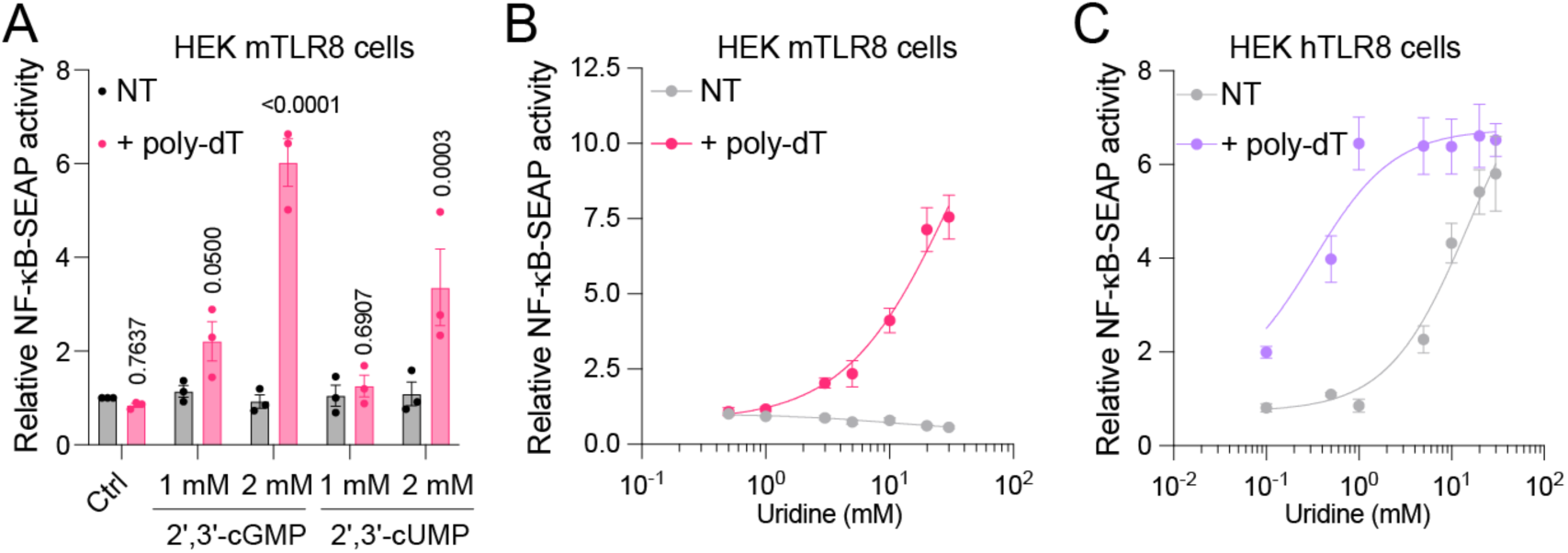
mTLR8 and hTLR8 responses to cyclic nucleotides and uridine. (**A-B**) HEK mTLR8 reporter cells were incubated overnight with (**A**) indicated combinations of 2′,3′-cGMP and 2′,3′-cUMP, alone and/or in combination with 17-mer poly-dT (5 μM), as indicated, (**B**) various doses of uridine, as indicated, alone or in combination with poly-dT (5 μM). (**C**) HEK hTLR8 reporter cells were incubated overnight with various doses of uridine, as indicated, alone or in combination with poly-dT (5 μM). (**A-C**) Optical density was assessed the next day. Unstimulated cells (Ctrl) served as negative control. Data are mean of *n* = 3 (A, C) or *n* = 4 (B) independent experiments and are shown as fold change of the optical density normalized to unstimulated cells (Ctrl) (± s.e.m. and [A] two-way ANOVA with uncorrected Fisher’s LSD tests shown comparing conditions of treatments with or without poly-dT).

**Supplementary Figure 3.**
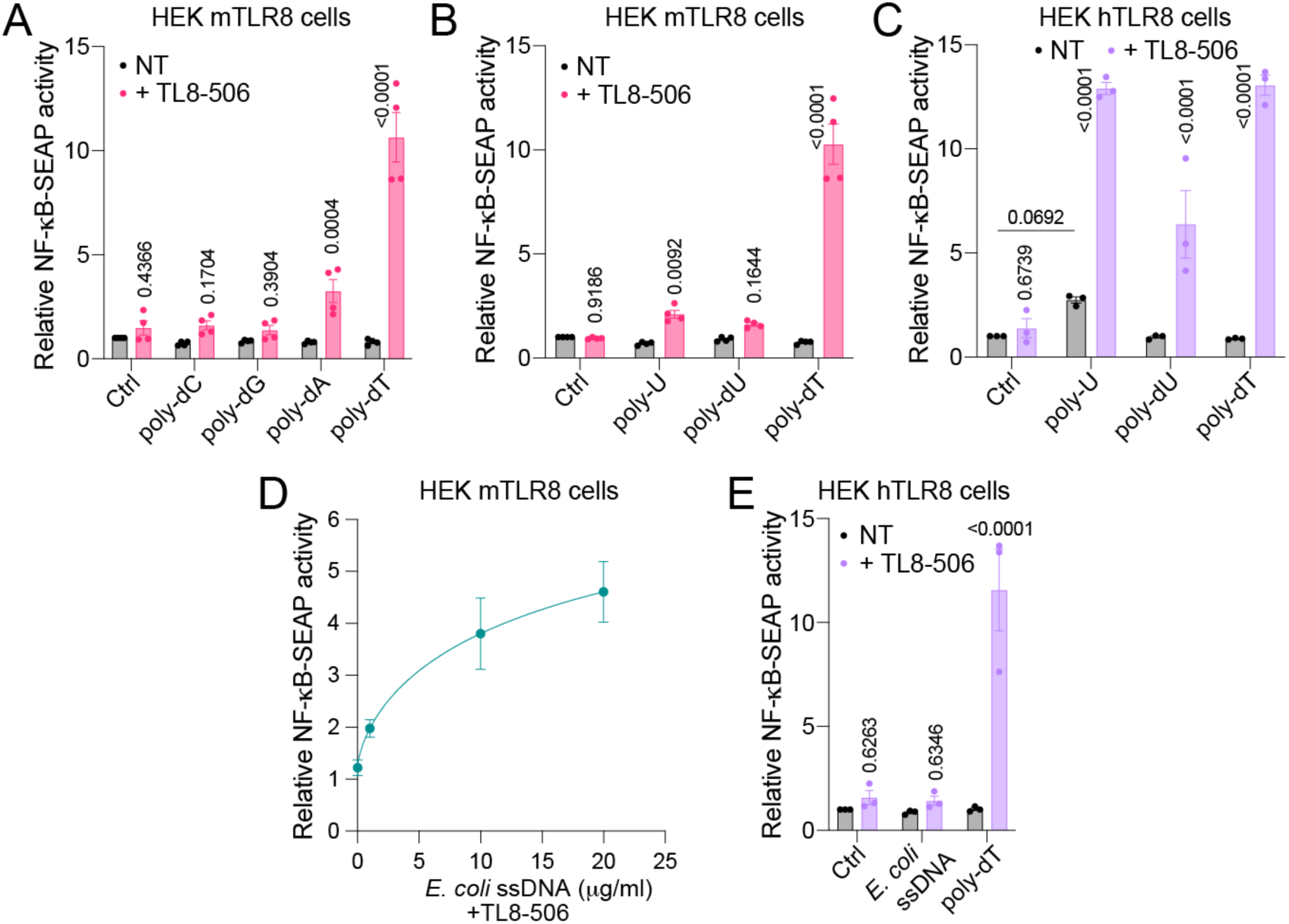
Site 2 agonists enhance TL8-506-induced mTLR8 and hTLR8 activation. HEK mTLR8 (**A, B**) or HEK hTLR8 (**C**) reporter cells were incubated overnight with poly-dA (10 μM), poly-dC (10 μM), poly-dG (10 μM), poly-dT (10 μM), poly-dU or poly-U (10 μg/ml each), as indicated, alone or in combination with TL8-506 (500 ng/ml). HEK (**D**) mTLR8 or (**E**) hTLR8 reporter cells were treated overnight with various doses of ssDNA purified from *E. coli,* as indicated, combined with TL8- 506 (100 ng/ml). (**A-E**) Optical density was assessed the next day. Unstimulated cells (Ctrl) served as negative control. Data are mean of *n* = 3 (C, E) or *n* = 4 (A, B, D) independent experiments. Data are shown as fold change of the optical density normalized to unstimulated cells (Ctrl) (± s.e.m. and [A-C, E] two-way ANOVA with uncorrected Fisher’s LSD tests shown comparing conditions of treatments with or without poly-dT or otherwise indicated pairs of conditions).

**Supplementary Figure 4.**
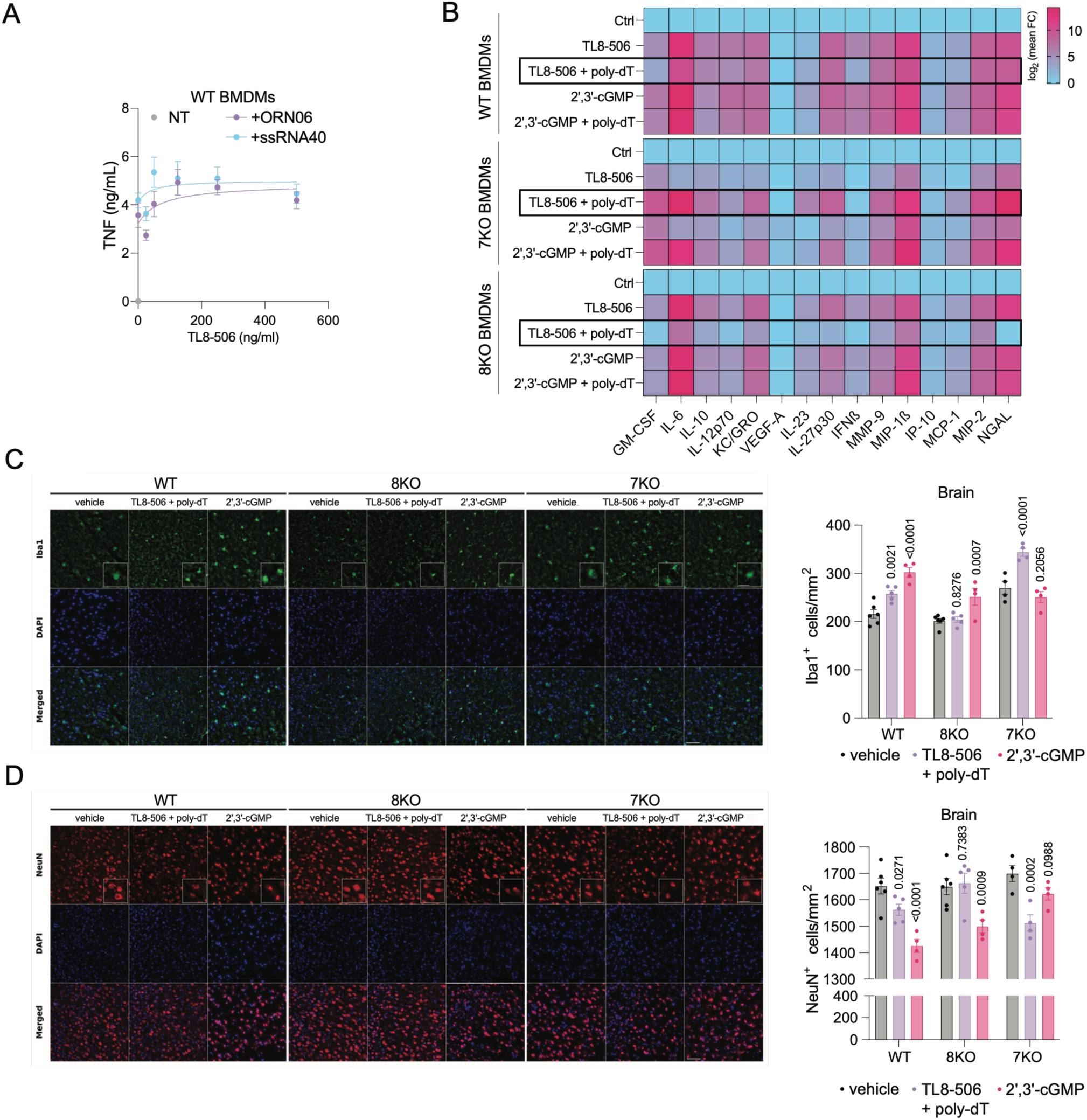
Inflammatory effects of mTLR8 agonists *in vitro* and *in vivo*. (**A**) BMDMs isolated from C57BL/6 mice (wild-type, WT) were incubated overnight with various doses of TL8-506, as indicated, alone or in combination with 20-mer poly-dT (5 μM), transfected ORN06 (10 μg/ml), or transfected ssRNA40^PS^ (10 μg/ml). Subsequently, TNF concentration levels in the supernatant were assessed by ELISA. Unstimulated cells (NT) served as negative control. Data are represented as mean ± s.e.m. (*n* = 4). (**B**) BMDMs isolated from WT, *Tlr7*^-/-^ (7KO), or *Tlr8*^-/-^ (8KO) mice were incubated overnight with 500 ng/ml TL8-506 or 10 mM 2′,3′-cGMP, with or without 20-mer poly-dT (5 µM). Cytokine concentrations were determined using a custom U-PLEX multiplex immunoassay (see *Methods*). Heatmap shows color-coded Log_2_-fold changes of cytokine production relative to unstimulated cells (Ctrl) for each genotype. (**C, D**) Intrathecal injection of the combination 1 μg of TL8-506 plus 0.8 nmol of poly-dT, 276 µg of 2′,3′-cGMP, or PBS (vehicle control) into WT (TL8-506 + poly-dT, *n* = 5; 2′,3′- cGMP, *n* = 4; PBS, *n* = 6), 8KO (TL8-506 + poly-dT, *n* = 5; 2′,3′-cGMP, *n* = 4; PBS, *n* = 6), or 7KO (TL8-506 + poly-dT, *n* = 4; 2′,3′-cGMP, *n* = 4; PBS, *n* = 4) mice. After 3 d, brain sections were immunostained using (**C**) Iba1 and (**D**) NeuN antibody (**left panels**, scale bar, 50 µm), and microglia and neurons, respectively, in the cerebral cortex were quantified (**right panels**). Data are shown as average ± s.e.m. and two-way ANOVA with uncorrected Fisher’s LSD tests shown comparing treatments with vehicle group for each genotype.

**Supplementary Figure 5.**
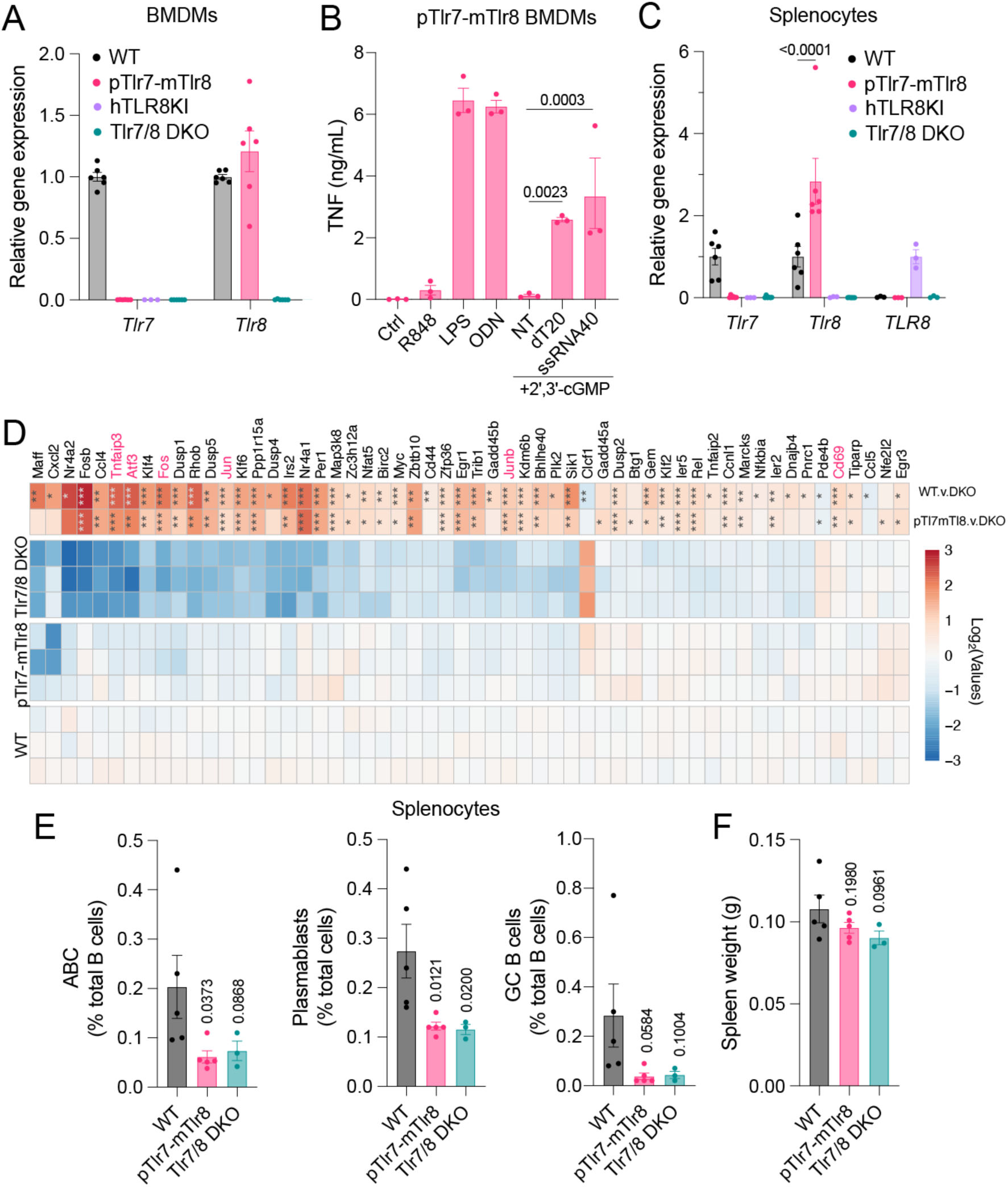
**Comparative characterization of pTlr7-mTlr8, hTLR8KI, and Tlr7/8 DKO mice**. (**A**) RT-qPCR analyses of total BMDM RNA from 6 C57BL/6 (wild-type, WT), 6 pTlr7-mTlr8, 3 hTLR8KI, and 6 Tlr7/8 DKO mice using primers directed against *mTlr7* or *mTlr8,* as indicated. (**B**) pTlr7- Tlr8 BMDMs were incubated overnight with 2′,3′-cGMP (10 mM), alone or in combination with 20-mer poly-dT (3 μM), or transfected ssRNA40^PS^ (10 μg/ml), as indicated. Unstimulated cells (Ctrl) and R848 (1 μg/ml) served as negative control, while LPS (100 ng/ml) and ODN1826 (500 nM) served as positive control. TNF concentrations in the supernatant were assessed by TNF ELISA the next day. (**C**) RT- qPCR analyses of total splenocytes from 6 WT, 6 pTlr7-mTlr8, 3 hTLR8KI, and 6 Tlr7/8 DKO mice using primers directed against *mTlr7, mTlr8, or hTLR8,* relative to *m18S*, as indicated. (**D**) Bulk RNA-seq of unstimulated splenocytes isolated from 3 WT, 3 pTlr7-mTlr8, and 3 Tlr7/8 DKO female mice. Heat map showing relative expression and log_2_ fold changes of select genes (rows) across splenocyte samples (columns). The 56 genes from the MSigDB Hallmark *TNFα signalling via NFκB* gene set that were significantly differentially expressed in one or both splenocyte comparisons are shown, with the relevant differential expression analyses indicated in the rightmost columns. Sample lines display log_2_ CPM gene expression values for each sample, relative to the average expression for WT splenocytes. The top two lines display log_2_ fold changes in gene expression between specific genotype groups as indicated by labels. Asterisks indicate genes significantly different between comparisons: * = *p* < 0.05, ** = *p* < 0.01, *** = *p* < 0.001. Colour scale (used for both sample expression and fold change values) was truncated at ±3. (**E**) Fractions of atypical B cells (ABCs, left, B220^+^CD19^+^CD21/35^-^CD23^-^CD11c^+^), plasmablasts (middle, CD3-CD11b-CD98+CD138hi), and GC B cells (right, B220^+^CD19^+^CD95^+^Bcl6^+^) from WT, pTlr7- mTlr8, and Tlr7/8 DKO mouse spleens, as assessed by flow cytometry. (**F**) Weight of spleens used in **E**. (**A, C**) Data are shown relative to WT mice (*mTlr7* and *mTlr8*) or hTLR8KI mice (*hTLR8*) (± s.e.m. and [**C**] two-way ANOVA with uncorrected Fisher’s LSD test shown). (**B**) Data are presented as mean of *n* = 3 mice (± s.e.m. and one-way ANOVA with uncorrected Fisher’s LSD tests shown compared to “2′,3′-cGMP + NT” condition) (**E**) Proportions of indicated cell subsets relative to total B cells counts for each spleen or (**F**) averaged total spleen weight are shown (± s.e.m. and one-way ANOVA with uncorrected Fisher’s LSD tests shown comparing to WT mice).

**Supplementary Figure 6.**
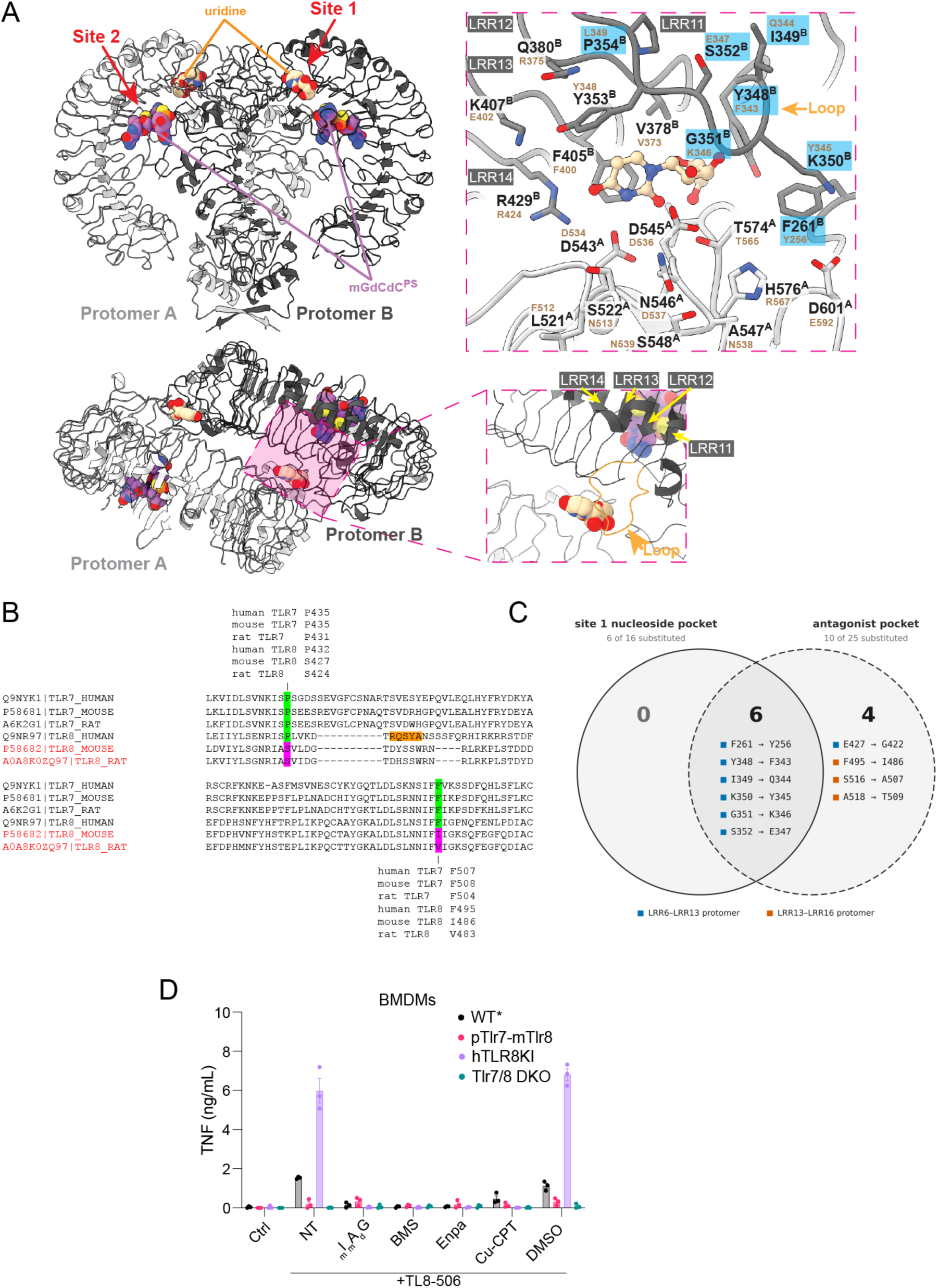
Analyses of mTlr7 and mTlr8 antagonistic sites. (**A**, top left) Crystal structure of hTLR8 dimer in complex with uridine at binding site 1 and 38-2 (_m_G_d_C_d_C^PS^) at binding site 2. Two TLR8 protomer ribbons are colored in light and dark grey, respectively. The uridine and _m_G_d_C_d_C^PS^ ligands are shown in VdW ball representation at each binding site. (**A**, bottom left) Alternative view of hTLR8-U-mGdCdC^PS^ at the protein dimerization interface with zoomed-in panel to show the location of leucine-rich-repeats (LRR11-LRR14) at the site 1. The LLR11-12 loop potentially modulating site 1 function is indicated with an orange arrow and colored in orange. (**A**, right) Agonist binding site (site 1) in hTLR8. Two TLR8 protomer ribbons are colored in light and dark grey, respectively. The protein residue side chains are shown for the residues either in contact with agonistic ligands or in difference with mTLR8 (mTLR8 residues are shown in small brown font). Residues highlighted in light blue are overlapping the antagonistic site. The ligand uridine is in ball-and-chain representation. The LRR11-12 loop is indicated by orange arrow. (**B**) Clustalw alignment of portions of hTLR7 (Q9NYK1), mTLR7 (P58681), rat TLR7 (A6K2G1), hTLR8 (Q9NR97), mTLR8 (P58682), and rat TLR8 (A0A8K0ZQ97) highlighting the amino acid variations at key residues of the antagonist binding site. The RQSYA motif previously thought to modulate mTLR8 function is highlighted in orange. (**C**) All residues substituted in the mTLR8 site 1 pocket also contribute to the antagonist-binding pocket. Non-conserved human-to- mouse TLR8 substitutions at residues contacting the site 1 nucleoside or antagonist are shown. Entries list the human residue and aligned mouse counterpart. Squares denote the LRR region of the protomer contributing each residue to the dimeric antagonist pocket. Conserved residues are omitted. (**D**) BMDMs from wild-type (WT), pTlr7-mTlr8, hTLR8KI, and Tlr7/8 DKO mice were pre-treated for 30 min with 5 μM _m_I_m_A_d_G^PS^ or 1 μM BMS-905, Enpatoran, or Cu-CPT-8m (CU-CPT) prior to overnight stimulation with TL8-506 at 50 ng/mL (hTLR8KI) or 1 μg/ml (other genotypes). mTNF levels in cell supernatants were quantified by ELISA. Data are presented as mean of *n* = 3 mice ± s.e.m.

**Supplementary Figure 7.**
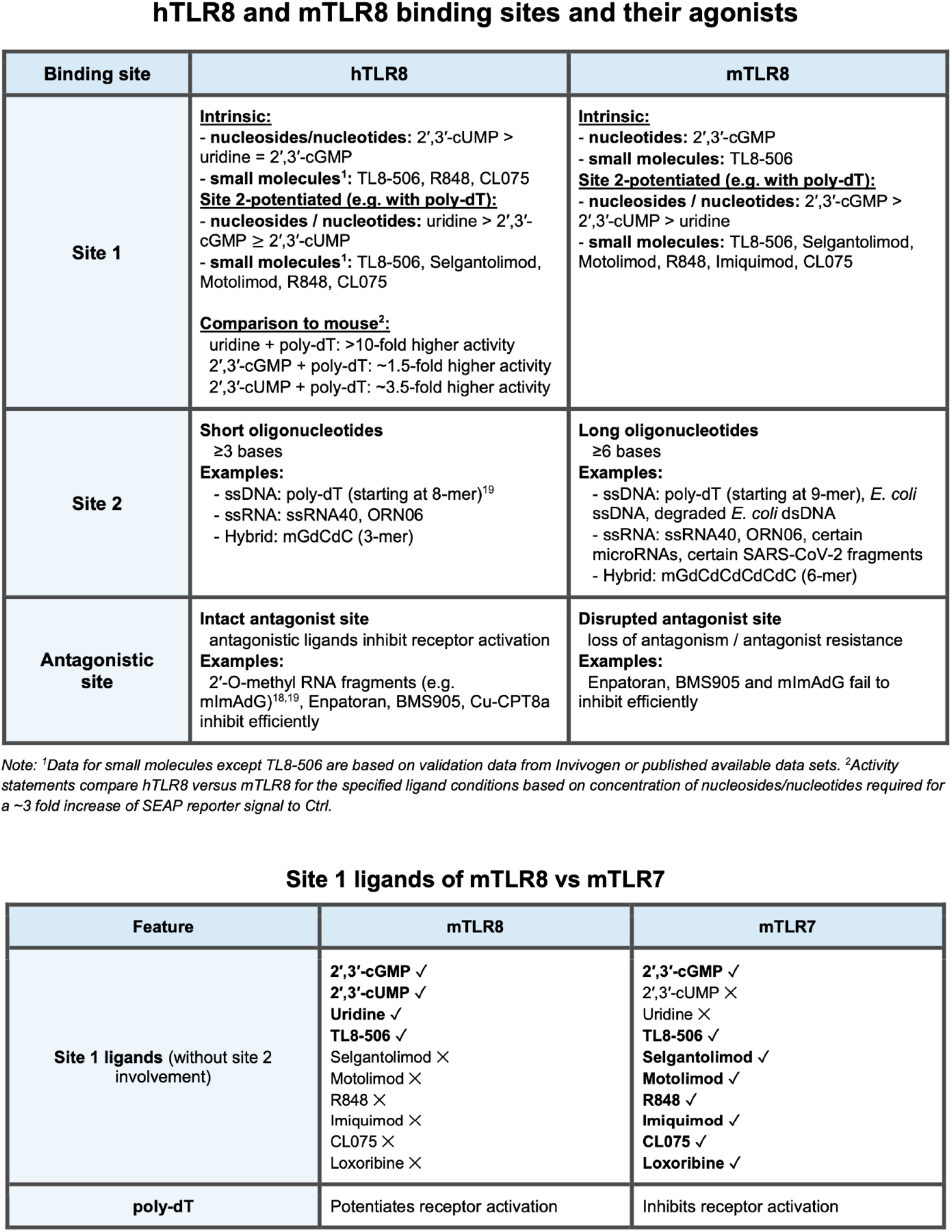
Summary of site activities for hTLR8, mTLR8, and mTLR7.

**Supplementary Figure 8.**
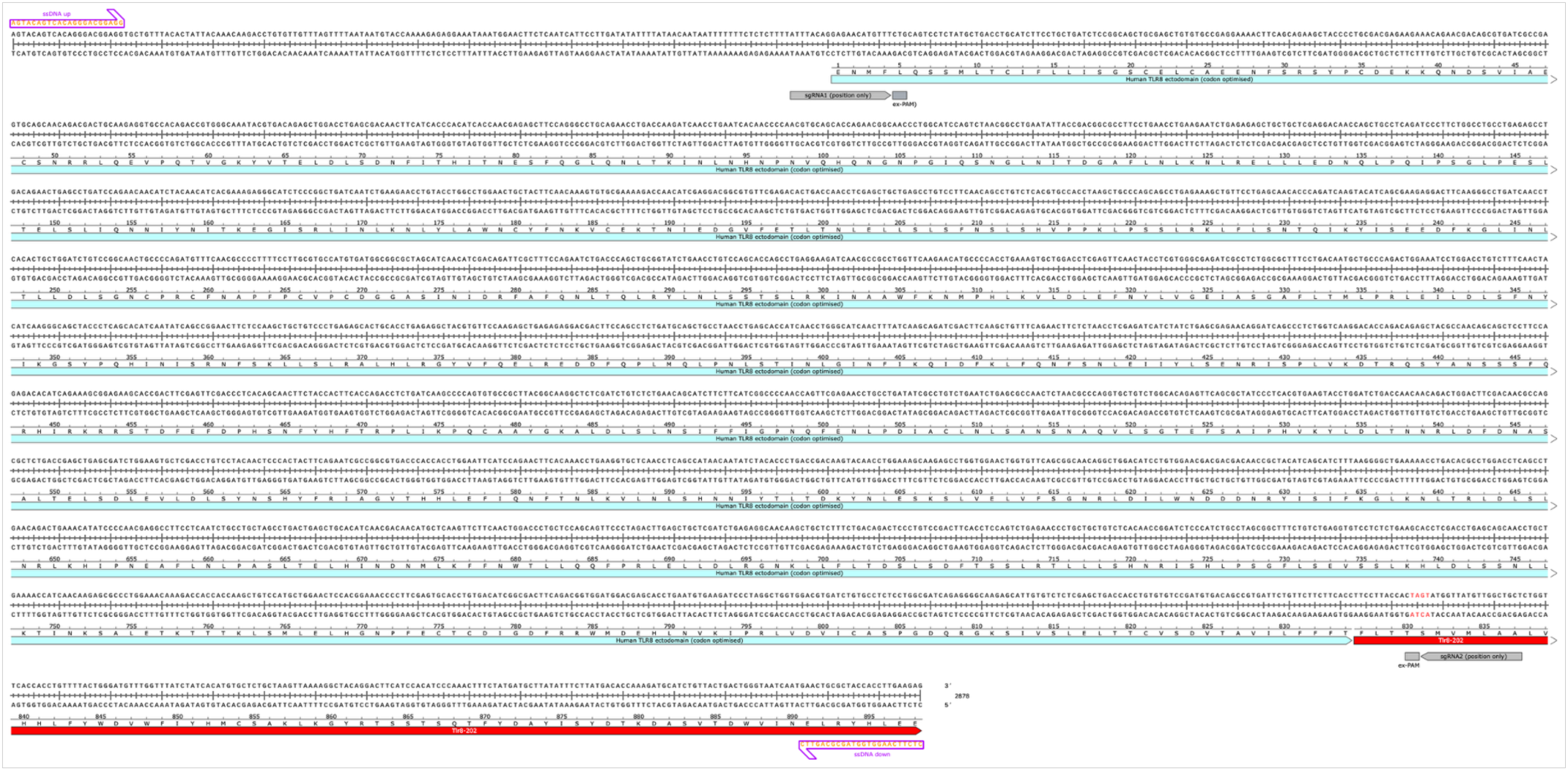
Repair template used to generate hTLR8KI mice. The sequence provided shows the entire repair template used, including 5’ and 3’ arms and the human TLR8 (mouse codon optimized) ectodomain (blue) which was used together with sgRNA1 and sgRNA2 to generate the hTLR8KI mice. The red sequence is that of the mTLR8 protein.

**Supplementary Figure 9.**
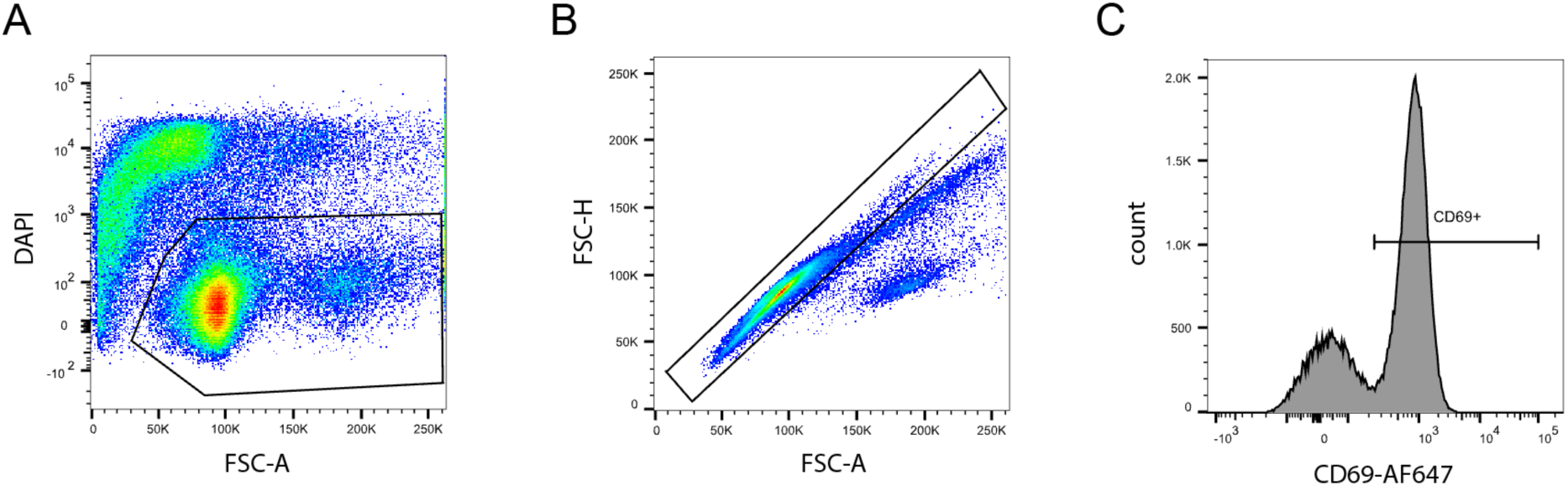
CD69^+^ splenocyte gating strategies. (**A-C**) Single-cell suspensions were analysed on a BD LSR-Fortessa X-20 flow cytometer using BD FACSDiva software, with 50,000 events (single, live cells) acquired per sample. Data analysis was subsequently conducted using FlowJo v10.9. Cell populations were first gated for live cells identified as the V450 (DAPI) negative population (**A**), followed by gating for single cells using forward scatter area (FSC-A) versus forward scatter height (FSC-H) parameters (**B**). Finally, CD69^+^ cells were gated within the live, single-cell populations (**C**). The representative sample shown is pTlr7-mTlr8 mouse splenocytes stimulated with the mouse TLR9 agonist ODN 1826, used as a positive control.

## References

1. Medzhitov, R. (2001). Toll-like receptors and innate immunity. Nat. Rev. Immunol. 1, 135–145. 10.1038/35100529.

2. Kawai, T., and Akira, S. (2010). The role of pattern-recognition receptors in innate immunity: update on Toll-like receptors. Nat. Immunol. 11, 373–384. 10.1038/ni.1863.

3. Janeway, C.A., and Medzhitov, R. (2002). Innate Immune Recognition. Annu. Rev. Immunol. 20, 197–216. 10.1146/annurev.immunol.20.083001.084359.

4. Nathan, C., and Cunningham-Bussel, A. (2013). Beyond oxidative stress: an immunologist’s guide to reactive oxygen species. Nat. Rev. Immunol. 13, 349–361. 10.1038/nri3423.

5. Lind, N.A., Rael, V.E., Pestal, K., Liu, B., and Barton, G.M. (2022). Regulation of the nucleic acid- sensing Toll-like receptors. Nat. Rev. Immunol. 22, 224–235. 10.1038/s41577-021-00577-0.

6. Heil, F., Hemmi, H., Hochrein, H., Ampenberger, F., Kirschning, C., Akira, S., Lipford, G., Wagner, H., and Bauer, S. (2004). Species-specific recognition of single-stranded RNA via toll-like receptor 7 and 8. Science 303, 1526–1529. 10.1126/science.1093620.

7. Diebold, S.S., Kaisho, T., Hemmi, H., Akira, S., and Reis e Sousa, C. (2004). Innate antiviral responses by means of TLR7-mediated recognition of single-stranded RNA. Science 303, 1529– 1531. 10.1126/science.1093616.

8. Fabbri, M., Paone, A., Calore, F., Galli, R., Gaudio, E., Santhanam, R., Lovat, F., Fadda, P., Mao, C., Nuovo, G.J., et al. (2012). MicroRNAs bind to Toll-like receptors to induce prometastatic inflammatory response. Proc Natl Acad Sci USA 109, E2110–6. 10.1073/pnas.1209414109.

9. Alexopoulou, L., Holt, A.C., Medzhitov, R., and Flavell, R.A. (2001). Recognition of double-stranded RNA and activation of NF-κB by Toll-like receptor 3. Nature 413, 732–738. 10.1038/35099560.

10. Jurk, M., Heil, F., Vollmer, J., Schetter, C., Krieg, A.M., Wagner, H., Lipford, G., and Bauer, S. (2002). Human TLR7 or TLR8 independently confer responsiveness to the antiviral compound R- 848. Nat. Immunol. 3, 499–499. 10.1038/ni0602-499.

11. Hemmi, H., Takeuchi, O., Kawai, T., Kaisho, T., Sato, S., Sanjo, H., Matsumoto, M., Hoshino, K., Wagner, H., Takeda, K., et al. (2000). A Toll-like receptor recognizes bacterial DNA. Nature 408, 740–745. 10.1038/35047123.

12. Oldenburg, M., Krüger, A., Ferstl, R., Kaufmann, A., Nees, G., Sigmund, A., Bathke, B., Lauterbach, H., Suter, M., Dreher, S., et al. (2012). TLR13 Recognizes Bacterial 23 *S* rRNA Devoid of Erythromycin Resistance–Forming Modification. Science 337, 1111–1115. 10.1126/science.1220363.

13. Roach, J.C., Glusman, G., Rowen, L., Kaur, A., Purcell, M.K., Smith, K.D., Hood, L.E., and Aderem, A. (2005). The evolution of vertebrate Toll-like receptors. Proc. Natl. Acad. Sci. 102, 9577–9582. 10.1073/pnas.0502272102.

14. Shibata, T., Ohto, U., Nomura, S., Kibata, K., Motoi, Y., Zhang, Y., Murakami, Y., Fukui, R., Ishimoto, T., Sano, S., et al. (2016). Guanosine and its modified derivatives are endogenous ligands for TLR7. Int. Immunol. 28, 211–222. 10.1093/intimm/dxv062.

15. Zhang, Z., Ohto, U., Shibata, T., Krayukhina, E., Taoka, M., Yamauchi, Y., Tanji, H., Isobe, T., Uchiyama, S., Miyake, K., et al. (2016). Structural Analysis Reveals that Toll-like Receptor 7 Is a Dual Receptor for Guanosine and Single-Stranded RNA. Immunity 45, 737–748. 10.1016/j.immuni.2016.09.011.

16. Tanji, H., Ohto, U., Shibata, T., Miyake, K., and Shimizu, T. (2013). Structural reorganization of the Toll-like receptor 8 dimer induced by agonistic ligands. Science 339, 1426–1429. 10.1126/science.1229159.

17. Zhang, Z., Ohto, U., Shibata, T., Taoka, M., Yamauchi, Y., Sato, R., Shukla, N.M., David, S.A., Isobe, T., Miyake, K., et al. (2018). Structural Analyses of Toll-like Receptor 7 Reveal Detailed RNA Sequence Specificity and Recognition Mechanism of Agonistic Ligands. Cell Rep. 25, 3371–3381.e5. 10.1016/j.celrep.2018.11.081.

18. Tanji, H., Ohto, U., Shibata, T., Taoka, M., Yamauchi, Y., Isobe, T., Miyake, K., and Shimizu, T. (2015). Toll-like receptor 8 senses degradation products of single-stranded RNA. Nat. Struct. Mol. Biol. 22, 109–115. 10.1038/nsmb.2943.

19. Sapkota, S., Zhang, Z., Wolf, C., Fujimura, A., Wenholz, D.S., Speir, M., Wilkinson-White, L., Chan, C., Percier, P., Ladaycia, A., et al. (2026). Defining the Immunomodulatory Determinants of 3-mer Oligonucleotides on TLR7 and TLR8 sensing. Preprint at Immunology, 10.64898/2026.04.29.720527.

20. Alharbi, A.S., Sapkota, S., Zhang, Z., Jin, R., Rupasinghe, E., Jayasekara, W.S.N., Yu, D., Speir, M., Wilkinson-White, L., Cubeddu, L., et al. (2026). 2′-O-Methyl-guanosine RNA fragments antagonize TLR7 and TLR8 to limit autoimmunity. Nat. Immunol. 27, 762–775. 10.1038/s41590-026-02429-2.

21. Forsbach, A., Nemorin, J.-G., Montino, C., Müller, C., Samulowitz, U., Vicari, A.P., Jurk, M., Mutwiri, G.K., Krieg, A.M., Lipford, G.B., et al. (2008). Identification of RNA sequence motifs stimulating sequence-specific TLR8-dependent immune responses. J. Immunol. Baltim. Md 1950 *180*, 3729– 3738. 10.4049/jimmunol.180.6.3729.

22. Hemmi, H., Kaisho, T., Takeuchi, O., Sato, S., Sanjo, H., Hoshino, K., Horiuchi, T., Tomizawa, H., Takeda, K., and Akira, S. (2002). Small anti-viral compounds activate immune cells via the TLR7 MyD88–dependent signaling pathway. Nat. Immunol. 3, 196–200. 10.1038/ni758.

23. Liu, J., Xu, C., Hsu, L.-C., Luo, Y., Xiang, R., and Chuang, T.-H. (2010). A five-amino-acid motif in the undefined region of the TLR8 ectodomain is required for species-specific ligand recognition. Mol. Immunol. 47, 1083–1090. 10.1016/j.molimm.2009.11.003.

24. Krüger, A., Oldenburg, M., Chebrolu, C., Beisser, D., Kolter, J., Sigmund, A.M., Steinmann, J., Schäfer, S., Hochrein, H., Rahmann, S., et al. (2015). Human TLR8 senses UR/URR motifs in bacterial and mitochondrial RNA. EMBO Rep. 16, 1656–1663. 10.15252/embr.201540861.

25. Ma, Y., Li, J., Chiu, I., Wang, Y., Sloane, J.A., Lü, J., Kosaras, B., Sidman, R.L., Volpe, J.J., and Vartanian, T. (2006). Toll-like receptor 8 functions as a negative regulator of neurite outgrowth and inducer of neuronal apoptosis. J. Cell Biol. 175, 209–215. 10.1083/jcb.200606016.

26. Demaria, O., Pagni, P.P., Traub, S., Gassart, A. de, Branzk, N., Murphy, A.J., Valenzuela, D.M., Yancopoulos, G.D., Flavell, R.A., and Alexopoulou, L. (2010). TLR8 deficiency leads to autoimmunity in mice. J. Clin. Invest. 120, 3651–3662. 10.1172/JCI42081.

27. Zhang, Z.-J., Guo, J.-S., Li, S.-S., Wu, X.-B., Cao, D.-L., Jiang, B.-C., Jing, P.-B., Bai, X.-Q., Li, C.-H., Wu, Z.-H., et al. (2018). TLR8 and its endogenous ligand miR-21 contribute to neuropathic pain in murine DRG. J. Exp. Med. 215, 3019–3037. 10.1084/jem.20180800.

28. Rael, V.E., Yano, J.A., Huizar, J.P., Slayden, L.C., Weiss, M.A., Turcotte, E.A., Terry, J.M., Zuo, W., Thiffault, I., Pastinen, T., et al. (2024). Large-scale mutational analysis identifies UNC93B1 variants that drive TLR-mediated autoimmunity in mice and humans. J. Exp. Med. 221, e20232005. 10.1084/jem.20232005.

29. Desnues, B., Macedo, A.B., Roussel-Queval, A., Bonnardel, J., Henri, S., Demaria, O., and Alexopoulou, L. (2014). TLR8 on dendritic cells and TLR9 on B cells restrain TLR7-mediated spontaneous autoimmunity in C57BL/6 mice. Proc. Natl. Acad. Sci. U. S. A. 111, 1497–1502. 10.1073/pnas.1314121111.

30. Jurk, M., Kritzler, A., Schulte, B., Tluk, S., Schetter, C., Krieg, A.M., and Vollmer, J. (2006). Modulating responsiveness of human TLR7 and 8 to small molecule ligands with T-rich phosphorothiate oligodeoxynucleotides. Eur. J. Immunol. 36, 1815–1826. 10.1002/eji.200535806.

31. Gorden, K.K.B., Qiu, X.X., Binsfeld, C.C.A., Vasilakos, J.P., and Alkan, S.S. (2006). Cutting edge: activation of murine TLR8 by a combination of imidazoquinoline immune response modifiers and polyT oligodeoxynucleotides. J. Immunol. Baltim. Md 1950 *177*, 6584–6587. 10.4049/jimmunol.177.10.6584.

32. Gorden, K.K.B., Gorski, K.S., Gibson, S.J., Kedl, R.M., Kieper, W.C., Qiu, X., Tomai, M.A., Alkan, S.S., and Vasilakos, J.P. (2005). Synthetic TLR Agonists Reveal Functional Differences between Human TLR7 and TLR8. J. Immunol. 174, 1259–1268. 10.4049/jimmunol.174.3.1259.

33. Butchi, N.B., Pourciau, S., Du, M., Morgan, T.W., and Peterson, K.E. (2008). Analysis of the Neuroinflammatory Response to TLR7 Stimulation in the Brain: Comparison of Multiple TLR7 and/or TLR8 Agonists. J. Immunol. 180, 7604–7612. 10.4049/jimmunol.180.11.7604.

34. Gorden, K.K.B., Qiu, X., Battiste, J.J.L., Wightman, P.P.D., Vasilakos, J.P., and Alkan, S.S. (2006). Oligodeoxynucleotides differentially modulate activation of TLR7 and TLR8 by imidazoquinolines. J. Immunol. Baltim. Md 1950 *177*, 8164–8170. 10.4049/jimmunol.177.11.8164.

35. Lu, H., Dietsch, G.N., Matthews, M.-A.H., Yang, Y., Ghanekar, S., Inokuma, M., Suni, M., Maino, V.C., Henderson, K.E., Howbert, J.J., et al. (2012). VTX-2337 Is a Novel TLR8 Agonist That Activates NK Cells and Augments ADCC. Clin. Cancer Res. 18, 499–509. 10.1158/1078-0432.CCR-11-1625.

36. Köllisch, G., Solis, F.V., Obermann, H.-L., Eckert, J., Müller, T., Vierbuchen, T., Rickmeyer, T., Muche, S., Przyborski, J.M., Heine, H., et al. (2022). TLR8 is activated by 5’-methylthioinosine, a Plasmodium falciparum-derived intermediate of the purine salvage pathway. Cell Rep. 39, 110691. 10.1016/j.celrep.2022.110691.

37. Bérouti, M., Lammens, K., Heiss, M., Hansbauer, L., Bauernfried, S., Stöckl, J., Pinci, F., Piseddu, I., Greulich, W., Wang, M., et al. (2024). Lysosomal endonuclease RNase T2 and PLD exonucleases cooperatively generate RNA ligands for TLR7 activation. Immunity 57, 1482–1496.e8. 10.1016/j.immuni.2024.04.010.

38. Wallach, T., Wetzel, M., Dembny, P., Staszewski, O., Krüger, C., Buonfiglioli, A., Prinz, M., and Lehnardt, S. (2020). Identification of CNS Injury-Related microRNAs as Novel Toll-Like Receptor 7/8 Signaling Activators by Small RNA Sequencing. Cells 9, 186. 10.3390/cells9010186.

39. Wallach, T., Mossmann, Z.J., Szczepek, M., Wetzel, M., Machado, R., Raden, M., Miladi, M., Kleinau, G., Krüger, C., Dembny, P., et al. (2021). MicroRNA-100-5p and microRNA-298-5p released from apoptotic cortical neurons are endogenous Toll-like receptor 7/8 ligands that contribute to neurodegeneration. Mol. Neurodegener. 16, 80. 10.1186/s13024-021-00498-5.

40. The Tabula Muris Consortium, Overall coordination, Logistical coordination, Organ collection and processing, Library preparation and sequencing, Computational data analysis, Cell type annotation, Writing group, Supplemental text writing group, and Principal investigators (2018). Single-cell transcriptomics of 20 mouse organs creates a Tabula Muris. Nature 562, 367–372. 10.1038/s41586-018-0590-4.

41. Lehmann, S.M., Krüger, C., Park, B., Derkow, K., Rosenberger, K., Baumgart, J., Trimbuch, T., Eom, G., Hinz, M., Kaul, D., et al. (2012). An unconventional role for miRNA: let-7 activates Toll- like receptor 7 and causes neurodegeneration. Nat. Neurosci. 15, 827–835. 10.1038/nn.3113.

42. Lehnardt, S., Henneke, P., Lien, E., Kasper, D.L., Volpe, J.J., Bechmann, I., Nitsch, R., Weber, J.R., Golenbock, D.T., and Vartanian, T. (2006). A Mechanism for Neurodegeneration Induced by Group B Streptococci through Activation of the TLR2/MyD88 Pathway in Microglia. J. Immunol. 177, 583–592. 10.4049/jimmunol.177.1.583.

43. Rosenberger, K., Dembny, P., Derkow, K., Engel, O., Krüger, C., Wolf, S.A., Kettenmann, H., Schott, E., Meisel, A., and Lehnardt, S. (2015). Intrathecal heat shock protein 60 mediates neurodegeneration and demyelination in the CNS through a TLR4- and MyD88-dependent pathway. Mol. Neurodegener. 10, 5. 10.1186/s13024-015-0003-1.

44. Zhang, W., Xiao, D., Mao, Q., and Xia, H. (2023). Role of neuroinflammation in neurodegeneration development. Signal Transduct. Target. Ther. 8, 267. 10.1038/s41392-023-01486-5.

45. Soni, C., Wong, E.B., Domeier, P.P., Khan, T.N., Satoh, T., Akira, S., and Rahman, Z.S.M. (2014). B Cell–Intrinsic TLR7 Signaling Is Essential for the Development of Spontaneous Germinal Centers. J. Immunol. 193, 4400–4414. 10.4049/jimmunol.1401720.

46. Rubtsov, A.V., Rubtsova, K., Fischer, A., Meehan, R.T., Gillis, J.Z., Kappler, J.W., and Marrack, P. (2011). Toll-like receptor 7 (TLR7)–driven accumulation of a novel CD11c+ B-cell population is important for the development of autoimmunity. Blood 118, 1305–1315. 10.1182/blood-2011-01-331462.

47. Crow, Y.J., and Gantier, M.P. (2026). TLR7 mutations leading to enhanced TLR7 signaling in humans. J. Hum. Immun. 2, e20250266, jhi.20250266. 10.70962/jhi.20250266.

48. Vlach, J., Bender, A.T., Przetak, M., Pereira, A., Deshpande, A., Johnson, T.L., Reissig, S., Tzvetkov, E., Musil, D., Morse, N.T., et al. (2021). Discovery of M5049: A Novel Selective Toll-Like Receptor 7/8 Inhibitor for Treatment of Autoimmunity. J. Pharmacol. Exp. Ther. 376, 397–409. 10.1124/jpet.120.000275.

49. Sreekantha, R.K., Mussari, C.P., Dodd, D.S., Pasunoori, L., Hegde, S., Posy, S.L., Critton, D., Ruepp, S., Subramanian, M., Salter-Cid, L.M., et al. (2022). Identification of 2-Pyridinylindole-Based Dual Antagonists of Toll-like Receptors 7 and 8 (TLR7/8). ACS Med. Chem. Lett. 13, 812–818. 10.1021/acsmedchemlett.2c00049.

50. Bérouti, M., Wagner, M., Greulich, W., Piseddu, I., Gärtig, J., Hansbauer, L., Müller-Hermes, C., Heiss, M., Pichler, A., Tölke, A.J., et al. (2025). Pseudouridine RNA avoids immune detection through impaired endolysosomal processing and TLR engagement. Cell, S0092867425006191. 10.1016/j.cell.2025.05.032.

51. Aluri, J., Bach, A., Kaviany, S., Chiquetto Paracatu, L., Kitcharoensakkul, M., Walkiewicz, M.A., Putnam, C.D., Shinawi, M., Saucier, N., Rizzi, E.M., et al. (2021). Immunodeficiency and bone marrow failure with mosaic and germline TLR8 gain of function. Blood 137, 2450–2462. 10.1182/blood.2020009620.

52. Fejtkova, M., Sukova, M., Hlozkova, K., Skvarova Kramarzova, K., Rackova, M., Jakubec, D., Bakardjieva, M., Bloomfield, M., Klocperk, A., Parackova, Z., et al. (2022). TLR8/TLR7 dysregulation due to a novel TLR8 mutation causes severe autoimmune hemolytic anemia and autoinflammation in identical twins. Am. J. Hematol. 97, 338–351. 10.1002/ajh.26452.

53. Morand, E.F., Dall’Era, M., Sanchez-Guerrero, J., Pearson, D.R., Werth, V.P., Wenzel, J., Roy, S., Kleinmond, C., Gühring, H., Sgouroudi, E., et al. (2026). Enpatoran, a Toll-like receptor 7/8 inhibitor, in moderate-to-severe systemic lupus erythematosus: findings from Cohort B of a multicentre, international, double-blind, placebo-controlled dose-finding phase 2 trial. The Lancet 407, 1809– 1824. 10.1016/S0140-6736(26)00463-0.

54. Chakraborty, S., Ye, J., Wang, H., Sun, M., Zhang, Y., Sang, X., and Zhuang, Z. (2023). Application of toll-like receptors (TLRs) and their agonists in cancer vaccines and immunotherapy. Front. Immunol. 14, 1227833. 10.3389/fimmu.2023.1227833.

55. Kirtland, M.E., Tsitoura, D.C., Durham, S.R., and Shamji, M.H. (2020). Toll-Like Receptor Agonists as Adjuvants for Allergen Immunotherapy. Front. Immunol. 11, 599083. 10.3389/fimmu.2020.599083.

56. Guiducci, C., Gong, M., Cepika, A.-M., Xu, Z., Tripodo, C., Bennett, L., Crain, C., Quartier, P., Cush, J.J., Pascual, V., et al. (2013). RNA recognition by human TLR8 can lead to autoimmune inflammation. J. Exp. Med. 210, 2903–2919. 10.1084/jem.20131044.

57. Brown, G.J., Cañete, P.F., Wang, H., Medhavy, A., Bones, J., Roco, J.A., He, Y., Qin, Y., Cappello, J., Ellyard, J.I., et al. (2022). TLR7 gain-of-function genetic variation causes human lupus. Nature 605, 349–356. 10.1038/s41586-022-04642-z.

58. Yu, P., Lübben, W., Slomka, H., Gebler, J., Konert, M., Cai, C., Neubrandt, L., Prazeres da Costa, O., Paul, S., Dehnert, S., et al. (2012). Nucleic acid-sensing Toll-like receptors are essential for the control of endogenous retrovirus viremia and ERV-induced tumors. Immunity 37, 867–879. 10.1016/j.immuni.2012.07.018.

59. Tian, L., Su, S., Dong, X., Amann-Zalcenstein, D., Biben, C., Seidi, A., Hilton, D.J., Naik, S.H., and Ritchie, M.E. (2018). scPipe: A flexible R/Bioconductor preprocessing pipeline for single-cell RNA- sequencing data. PLOS Comput. Biol. 14, e1006361. 10.1371/journal.pcbi.1006361.

60. Liao, Y., Smyth, G.K., and Shi, W. (2019). The R package Rsubread is easier, faster, cheaper and better for alignment and quantification of RNA sequencing reads. Nucleic Acids Res. 47, e47–e47. 10.1093/nar/gkz114.

61. Durinck, S., Spellman, P.T., Birney, E., and Huber, W. (2009). Mapping identifiers for the integration of genomic datasets with the R/Bioconductor package biomaRt. Nat. Protoc. 4, 1184–1191. 10.1038/nprot.2009.97.

62. Robinson, M.D., McCarthy, D.J., and Smyth, G.K. (2010). edgeR : a Bioconductor package for differential expression analysis of digital gene expression data. Bioinformatics 26, 139–140. 10.1093/bioinformatics/btp616.

63. Robinson, M.D., and Oshlack, A. (2010). A scaling normalization method for differential expression analysis of RNA-seq data. Genome Biol. 11, R25. 10.1186/gb-2010-11-3-r25.

64. Law, C.W., Chen, Y., Shi, W., and Smyth, G.K. (2014). voom: precision weights unlock linear model analysis tools for RNA-seq read counts. Genome Biol. 15, R29. 10.1186/gb-2014-15-2-r29.

65. Ritchie, M.E., Phipson, B., Wu, D., Hu, Y., Law, C.W., Shi, W., and Smyth, G.K. (2015). limma powers differential expression analyses for RNA-sequencing and microarray studies. Nucleic Acids Res. 43, e47–e47. 10.1093/nar/gkv007.

66. Phipson, B., Lee, S., Majewski, I.J., Alexander, W.S., and Smyth, G.K. (2016). Robust hyperparameter estimation protects against hypervariable genes and improves power to detect differential expression. Ann. Appl. Stat. 10. 10.1214/16-AOAS920.

67. Liberzon, A., Birger, C., Thorvaldsdóttir, H., Ghandi, M., Mesirov, J.P., and Tamayo, P. (2015). The Molecular Signatures Database Hallmark Gene Set Collection. Cell Syst. 1, 417–425. 10.1016/j.cels.2015.12.004.

68. Subramanian, A., Tamayo, P., Mootha, V.K., Mukherjee, S., Ebert, B.L., Gillette, M.A., Paulovich, A., Pomeroy, S.L., Golub, T.R., Lander, E.S., et al. (2005). Gene set enrichment analysis: A knowledge-based approach for interpreting genome-wide expression profiles. Proc. Natl. Acad. Sci. 102, 15545–15550. 10.1073/pnas.0506580102.

